# Heterogeneity in heat shock response dynamics caused by translation fidelity decline and proteostasis collapse

**DOI:** 10.1101/822072

**Authors:** Nadia Vertti-Quintero, Simon Berger, Xavier Casadevall i Solvas, Cyril Statzer, Jillian Annis, Peter Ruppen, Stavros Stavrakis, Collin Y. Ewald, Rudiyanto Gunawan, Andrew deMello

## Abstract

Genetics, environment, and stochasticity influence the rate of ageing in living organisms. Individual *Caenorhabditis elegans* that are genetically identical and cultured in the same environment have different lifespans, suggesting a significant role of stochasticity in ageing. We have developed a novel microfluidic methodology to measure heat-shock response as a surrogate marker for heterogeneity associated with lifespan and have quantified the heat-shock response of *C. elegans* at the population, single individual, and tissue levels. We have further mathematically modelled our data to identify the major drivers determining such heterogeneity. This approach demonstrates that protein translation and degradation rate constants explain the individuality of the heat-shock time-course dynamic. We observed a decline of protein turnover capacity in early adulthood, co-incidentally occurring as the predicted proteostasis collapse. We identified a decline of intestinal response as the tissue that underlies the individual heterogeneity. Additionally, we verified that individuals with enhanced translation fidelity in early adulthood live longer. Altogether, our results reveal that the stochastic onset of proteostasis collapse of somatic tissues during early adulthood reflects individual protein translation capacity underlying heterogenic ageing of isogenic *C. elegans*.

## Introduction

Medical breakthroughs, such as antibiotics and vaccines, as well as improved health care have led to a doubling in human life expectancy in developed countries over the last 200 years [1]. The proportion of European population older than 65 years has been projected to exceed 20% by 2025, while in the US, the aged population is expected to double between 2010 and 2050 and reach 89 million [2]. However, the extension of lifespan has not necessarily been accompanied by a proportional increase in healthspan – the length of healthy life – and the final 20% of human lifespan is commonly associated with some degree of morbidity [1]. As such, a focal point in ageing research is the formulation of medical interventions that compress late-life morbidity [1–4]. One major challenge toward this aim is that the beneficial effects and efficacy of therapeutic treatments likely vary among individuals because of the heterogeneity of human ageing [2]. For instance, human monozygotic twins show different lengths in lifespan [5], discordance in the development of type I diabetes [6], and even major variability in the size of organs [7]. Although the genetic and environmental influences on ageing have been studied extensively, heterogeneity and stochasticity of the ageing process are much less well understood.

In this study, we investigated inter-individual heterogeneity in the ageing process using the model organism *Caenorhabditis elegans*. The nematode *C. elegans* is a small (*ca* 1 mm in length) and fast reproducing hermaphrodite that is particularly amenable to large scale studies because of the ease in maintaining a genetically identical (isogenic) population under tightly controlled conditions. Moreover, a typical lifespan of between three to four weeks makes the organism highly suitable for ageing research. By keeping the genetic background and environment uniform, any differences in the ageing process among a *C. elegans* population should thus result from non-genetic and other stochastic factors. Strikingly, isogenic *C. elegans* kept in the same environment and food source show large differences in lifespan, with some individuals dying on day 10, whilst others die on day 30 of adulthood [8–13]. Interventions that alter the lifespan of *C. elegans* have also been shown to affect the heterogeneity in the lifespan through a linear temporal scaling [14].

Besides lifespan, heterogeneity has been observed and studied in *C. elegans* for various phenotypes, such as cell fate [15], cell morphogenesis [16], synthetic lethality [17], microbiome-host interactions [18], body size [19], developmental speed [19], number of offspring [19], stress responses [20, 21], redox potential [22], and behaviour [23]. Of particular pertinence to the current study is the heterogeneity in the heat shock response (HSR), which has been tied to variability in lifespan, where a stronger HSR is indicative of longer lifespans among isogenic *C. elegans* [20]. Heat stresses are known to cause proteins to unfold, and unfolded proteins can further undergo entanglement and non-specific aggregation, limiting correct protein function [24]. Heat shock response is an ancient and highly conserved molecular mechanism that evolves as an adaptation and survival strategy to disruptions to cellular proteostasis [25]. The central element of the HSR is the feedback regulation where a heat shock factor (HSF) controls the expression of its deactivating molecule heat shock protein (HSP). HSPs are a family of proteins that protect the cells from the effects of stress, and many HSPs are chaperones that aid protein refolding [24, 25]. Under physiological conditions, HSFs are bound stoichiometrically to HSPs in the cytosol. Upon heat stress, increasing amount of cytosolic unfolded proteins recruits HSPs, thereby freeing up HSF. The released HSF trimerizes, translocates to the nucleus, and binds to its cognate heat shock element (HSE) site to activate the expression of HSPs. A higher expression of HSPs will continue until free cytosolic HSFs are again bound by HSPs.

In *C. elegans*, some types of HSPs, including *hsp-16.1, hsp-16.2, hsp-16.41, hsp-16.48* and *hsp-70*, are upregulated 16-fold within the first 30 minutes of a heat stress [26]. The expression of the *hsp-16.2* gene has previously been followed non-invasively in living *C. elegans* by genetically inserting an integrated transgene that expresses green fluorescent protein (GFP) under the control of the *hsp-16.2* promoter (P*hsp-16.2*∷GFP). Multiple studies using P*hsp-16.2*∷GFP *C. elegans* have shown that the expression of the GFP reporter predicts longevity of individuals [20, 27, 28]. This suggests that the capability of nematodes to respond to heat stress can be used as a predictive biomarker for their longevity. In our study, we took advantage of the relationship between HSR and lifespan in *C. elegans*, and used P*hsp-16.2*∷GFP to study the factor(s) contributing to ageing heterogeneity in an isogenic *C. elegans* population. Although the heterogeneity in *C. elegans* longevity has also been studied using other reporters, for example the oxidative response gene P*sod-3*∷GFP [8], splicing reporters [29], and P*mir-71*∷GFP and P*mir-246*∷GFP [9] microRNA promoters, we chose the P*hsp-16.2*∷GFP reporter for our study as it is a predictor of longevity as early as day-1 of adulthood [20].

To characterize HSR dynamics among individual in large *C. elegans* cohorts, we developed two novel microfluidic systems; one for screening fluorescent protein reporter expression in large numbers of individuals at specific time points (screening), and another for sorting the population according to the characteristics of such expression. We also employed a previously developed microfluidic device for high-resolution time-series GFP measurements of HSR dynamics in single *C. elegans* [30]. At the population level, we found that the heterogeneity in HSR within an isogenic TJ375 (P*hsp-16.2*∷GFP) [20] population generally follows a normal (Gaussian) distribution. Through sorting and screening experiments after heat-shock, we observed that isogenic *C. elegans* exhibit diverse HSR dynamics. We confirmed the heterogeneity in HSR dynamics by measuring the time-course of fluorescence in individual *C. elegans* TJ375 (P*hsp-16.2*∷GFP) after heat-shock. By applying mathematical modelling and parameter estimation to our HSR data, we identified the variability of protein translation and degradation rate constants as the underlying source of the population heterogeneity of HSR dynamics. We further confirmed (by lifespan assay) that faster HSR dynamics is concomitant with longer lifespan. Moreover, by characterizing the heterogeneity of HSR in *C. elegans* populations of two different ages (day-1 and day-2 adulthood), we observed a rapid age-associated decline in translation and degradation rate constants and thus a slowdown of HSR dynamics during early adulthood. Finally, looking at HSR dynamics within an individual *C. elegans*, we found that the centre region of the nematodes responds the quickest to heat-shock, and the HSR dynamics slow down towards the extremities of the animals.

## Results

### Time-course observation of heat shock response reveals deterministic heterogeneity in HSR at the population level of isogenic *C. elegans*

We used the transgenic nematode TJ375 (P*hsp-16.2*∷GFP) carrying a plasmid with GFP expression under the control of heat shock protein *hsp-16.2* promoter [20], for *in-vivo* assessments of HSR in *C. elegans* (Figure 1a,b). Consistent with previous reports [31], we found an agreement between the time-course of the endogenous HSP-16.2 and that of the GFP expression, via Western Blotting (Figure 1c,d, Supplementary Figure 1). Here, we developed a novel microfluidic system for measuring the expression of fluorescent protein reporters in *C. elegans* in a high-throughput and biocompatible manner (Figure 1e, Supplementary Figure 2), as detailed in Methods. We employed this microfluidic system to study the heterogeneity of HSR in an isogenic nematode population by GFP expression quantification of transgenic nematodes at multiple time points after heat stress. To the best of our knowledge, a time-course (dynamic) characterization of HSR for a constant nematode population upon heat-shock has not been previously performed.

**Figure 1.**
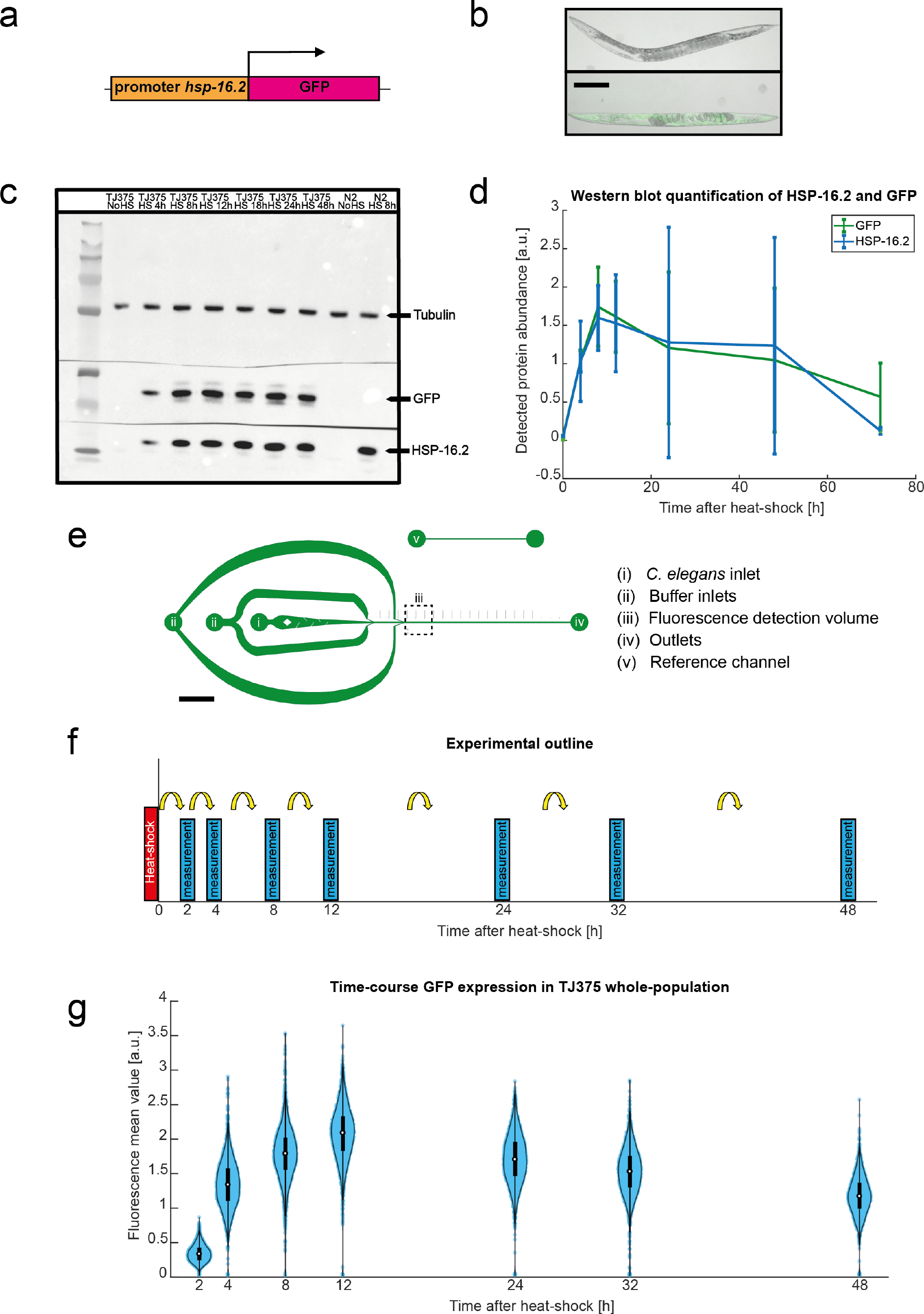
Longitudinal *C. elegans* experimental HSR characterization at population level. (a) Outline of the gene mutation of strain TJ375 [P*hsp-16.2*∷GFP(gpIs1)] [20]. (b) Bright field images of TJ375 animals (above) without heat-shock and (below) after heat-shock (fluorescent image overlaid to bright field). GFP is apparent on *C. elegans* hours after heat-shock. Bar scale represents 200 μm. (c) Image of a western blot for protein detection and quantification of HSP-16.2, GFP and Tubulin on *C. elegans* samples. Seven samples of the strain TJ375 were probed (from left to right: Without heat-shock and 4, 8, 12, 24, 48 and 72 hours after heat-shock), as well as 2 samples of “wild type” strain N2 (without heat-shock treatment and 8h after heat-shock). (d) Expression of endogenous GFP and HSP-16.2 of samples shown in (c), quantified with western blot. The error bars represent the standard deviation at each time point for 3 biological repeats. (e) Schematic of the microfluidic device for *C. elegans* fluorescent protein screening. The “*C. elegans* inlet” (i) incorporates wall-embedded PDMS blades to untangle animals as they enter the device. Additional buffer inlets (ii) enable efficient spacing of *C. elegans* in the main channel. (iii) Fluorescence detection volume where the optical detection system is focused onto. Animals exit via the device outlet (iv) and calibration is performed using the reference channel (v). The flow direction is indicated by the black arrow. Scale bars is 2.5 mm. (f) Experimental layout of *C. elegans* population screening experiments. (g) Violin plots of the fluorescence mean values of an isogenic population of *C. elegans* TJ375 (*n* = 2280) measured at determined time points (2, 4, 8, 12, 24, 32 and 48 hours) after heat-shock, as the experimental layout shown in (f). The thicker black lines inside the violins represent 50 percent of the population.

The details of the microfluidic system and the experimental procedure are described in Methods. Briefly, adult TJ375 *C. elegans* suspended in M9 buffer enter the microfluidic device through a main inlet and flow in a serial and non-overlapping manner through a main straight channel, supported by secondary buffer inlets (Figure 1e). The optical detection system integrates a detection volume, orthogonal to the direction of the flow stream. In this manner, when an animal passes through the detection volume, GFP expression (fluorescence intensity) along the body of individual *C. elegans* is recorded (Supplementary Figure 2). To assess the heterogeneity of HSR, we heat-shocked isogenic day-2 adult TJ375 *C. elegans* for 1-hour at 37° C. We then passed the nematodes through the microfluidic system and measured the whole-body GFP expression of the population at 2, 4, 8, 12, 24, 32 and 48 hours after the heat-shock (Figure 1f). We recorded the whole-body GFP fluorescence intensities of 2280 individual transgenic *C. elegans.* For all time points, the HSR heterogeneity in the nematode population follows a Gaussian distribution (Figure 1g and Supplementary Figure 3). The standard deviation of fluorescence intensities increases initially and then remains approximately constant beyond 4 hours after heat-shock (Supplementary Figure 3). This is in agreement with a previous study using the same transgenic nematodes [20]. The HSR heterogeneity does not differ markedly between TJ375 and a mutant with epigenetically suppressed HSR, LSD2088 (P*hsp-16.2*∷GFP, *jmjd-3.1(gk384)*) [32]) (Supplementary Figure 4).

The above cross-sectional measurements of the population give information on the heterogeneity of time-dependent GFP expression on a TJ375 *C. elegans* cohort, but do not specify the individual contributions from single animals. Thus, we are unable to determine whether the observed heterogeneity of HSR arises from differences in the individual HSR dynamics (*i.e.* each animal following a different HSR time-course) or from static variability (*i.e.* each animal following the same HSR time-course, but with the level of HSP or GFP expression varying between animals).

### Heterogeneity in the dynamics of HSR in isogenic *C. elegans* sub-populations is observed

To shed light on the source of the observed HSR heterogeneity among TJ375 *C. elegans*, we adapted the aforementioned microfluidic system to allow for the sorting of nematode populations into up to three groups according to their whole-body fluorescence signal in a high-throughput and biocompatible manner (Figure 2a, Supplementary Figure 2), as detailed in Methods. Using this microfluidic system, we sorted a day-2 adult TJ375 *C. elegans* population after a 1-hour 37° C heat-shock (see Figure 2b). First, the heat-shocked population was sorted into two groups based on the whole-body GFP fluorescence at 4 hours after heat shock: a *High* sub-population for nematodes with a whole-body fluorescence above the median level, and a *Low* sub-population for those with a whole-body fluorescence below the median level. Subsequently, at 8 hours after heat-shock, we repeated the sorting process for each sub-populations. For the *High* sub-population, we separated the top 16% from the bottom 84% of fluorescence intensities, and respectively labelled them as *High-High* and *High-Low* (see Figure 2c). Mirroring this sorting strategy, we sorted the *Low* sub-population at 8 hours after heat-shock to separate the bottom 16% from the top 84%, and respectively labelled them as *Low-Low* and *Low-High* (see Figure 2c). Separation in this manner, captures the upper (lower) tail of the *High* (*Low*) sub-population at one standard deviation above (below) the mean. We further screened GFP expression of the four resulting sub-populations at several time points after the sorting experiments – at 12, 25, and 33 hours after heat-shock (coloured in red, yellow, blue, and green; Figure 2c). By performing these double-sorting experiments, we were able to follow dynamically distinct sub-populations and understand better the heterogeneity of HSR dynamics in the TJ375 *C. elegans.*

**Figure 2.**
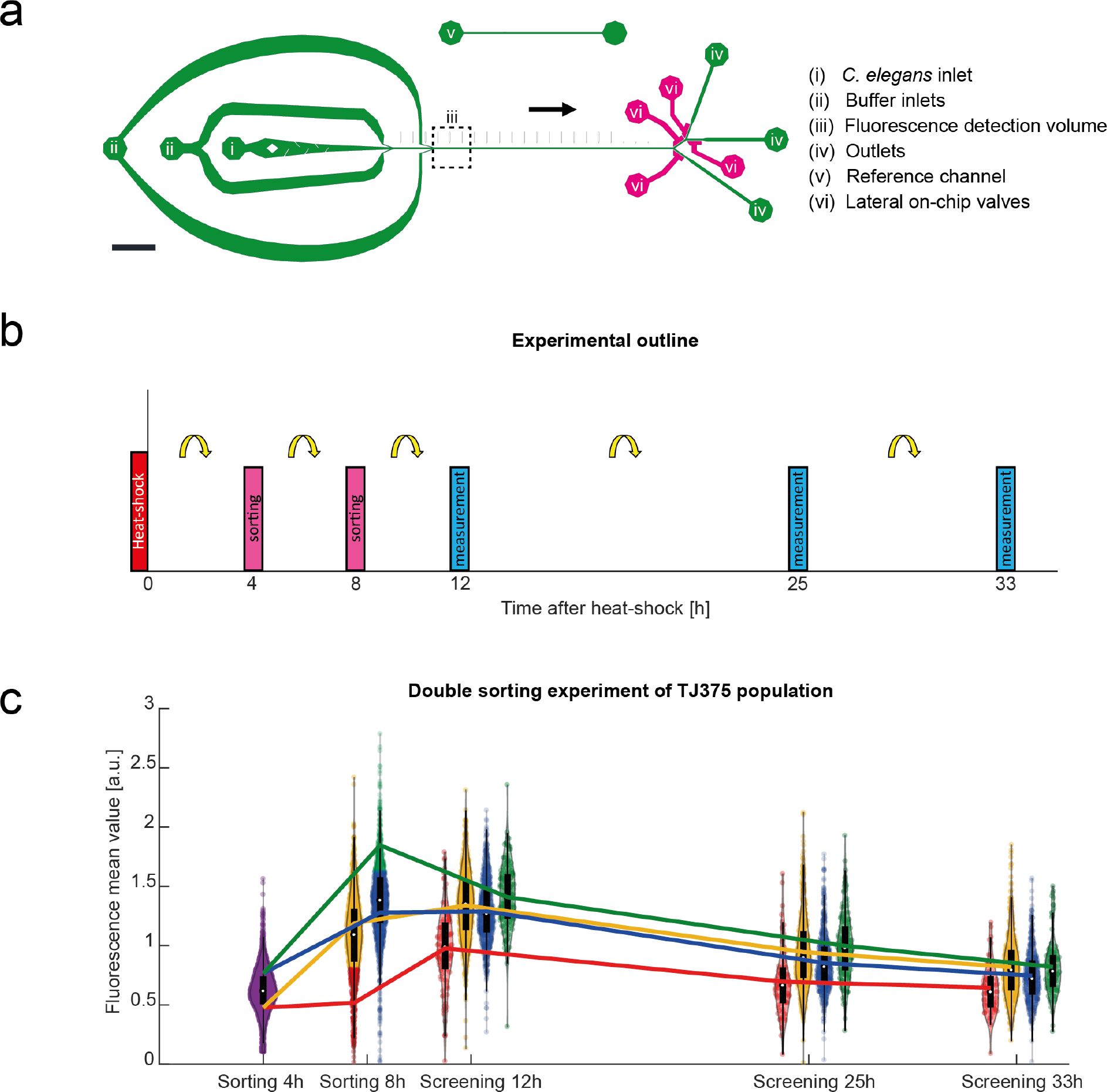
*C. elegans* experimental HSR characterization at sub-population level. (a) Schematic of the microfluidic device for *C. elegans* sorting. The fluidic layers are shown in green and the control layer in pink. The flow direction is indicated by the black arrow. Scale bar is 2.5 mm. The “*C. elegans* inlet” (i) incorporates wall-embedded PDMS blades to untangle animals as they enter the device. Additional buffer inlets (ii) enable efficient spacing of animals in the main channel. (iii) Fluorescence detection volume where the optical detection system is focused onto. Sorted animals exit via one of the device outlets (iv) and calibration is performed using the reference channel (v). Lateral on-chip valves are labelled (vi). (b) Experimental layout of *C. elegans* population sorting experiments. (c) Double sorting and subsequent GFP intensity measurement of a *C. elegans* population (*n* = 2600), following layout shown in (b), at different time points after heat-shock. We show each measured population with violin plots. Initial sorting was performed 4 hours after heat-shock, dividing the population into a top and bottom 50 %. A secondary sorting step (8 hours after heat-shock) further separated the top 50 % into its top 16 % and bottom 84 %. Similarly, the initial bottom 50 % of the population was further sorted into a bottom 16 % and a top 84 %. The four resultant groups were screened at posterior time points (12, 25 and 32 hours). The change in the mean value of each group at each sorting or screening time is shown with a distinct colour line per group, *i.e.* green: “*high-high*” group, blue: “*high-low*”, yellow: “*low-high*” and red: “*low-low*”.

We observed that within 4 hours of the first sorting (*i.e.* 8 hours after heat-shock), the whole-body GFP average fluorescence of each sub-population returned to an approximately Gaussian distribution (see Figure 2c). Comparing the *High* and *Low* sub-populations at 8 hours after heat-shock, we also noted a large overlap between their whole-body fluorescence average distributions. That said, the *High* sub-population maintained a statistically significant higher average whole-body fluorescence signal than the *Low* sub-population (two sample *t*-test *p*-value < 1 × 10^−5^). Among the four sub-populations resulting from the second sorting experiment, the *High-High* sub-population – the strongest responders to heat-shock – maintained the highest average fluorescence over the remaining duration of the experiment (green line; Figure 2c), having a maximum recorded fluorescence at 8 hours after heat-shock; whilst the *Low-Low* sub-population – the weakest responders – had the lowest average fluorescence (red line; Figure 2c), with a maximum recorded GFP expression at 12 hours after heat-shock. The *High*-*Low* and *Low-High* sub-populations displayed strikingly similar HSR dynamics post-sorting. But interestingly, the *Low-High* group had a slightly (but statistically significant) higher average fluorescence than the *High-Low* group at and beyond 12 hours after heat-shock (two sample *t*-test *p*-value = 3.99 × 10^−4^; yellow and blue lines in Figure 2c, Supplementary Table 1). The cross-over in the average fluorescence between the *High-Low* and *Low-High*, as well as between *High-High* and *High-Low*, suggest that a number of animals have a slow response to heat-shock. Finally, differences among the four sub-populations were observed to diminish over time. The results from these double-sorting experiments support the notion that the heterogeneity of HSR in TJ375 *C. elegans* is a consequence of variability in the HSR dynamics among the animals, *i.e.* each nematode follows a different time-course HSR trajectory.

### Time-lapse imaging reveals heterogeneity in the dynamics of HSR in individual *C. elegans*

Our initial sorting-screening experiments suggest a significant animal-to-animal heterogeneity in HSR dynamics, even within an isogenic population that is kept in the same environment. To understand better the key molecular mechanisms that give rise to this heterogeneity, we obtained a longitudinal observation of the HSR time-course by measuring GFP fluorescence in single TJ375 *C. elegans* using time-lapse microscopy. For this purpose, we adapted a microfluidic platform for long-term and high-resolution imaging of immobilized *C. elegans* that was previously developed in our laboratory [30] (Figure 3a-b; details on the device in Methods section). We confirmed that the device itself does not induce HSR response in TJ375 *C. elegans* for the same duration of the experiment (Supplementary Figure 5a). We characterized the HSR dynamics after 1-hour 37° C heat-shock for fifteen day-2 adult TJ375 *C. elegans* (Figure 3c, Supplementary Figure 5b, Supplementary Video 1), measuring the GFP expression intensity hourly over 48 hours. The results of this experiment are presented in Figure 3d, where the different coloured lines illustrate the whole-body fluorescence of individual *C. elegans* as well as the average GFP emission across the 15 *C. elegans* (bold black line). In general, GFP expression shows a sharp increase after heat-shock, peaking between 6 and 24 hours after heat shock and then decreasing following a typical exponential decay dynamic. However, it is noticeable that the rate of the GFP expression increase following heat-shock differs between individuals. Accordingly, the longitudinal time-course measurements for single animals confirm a significant animal-to-animal heterogeneity in the HSR dynamics, as suggested by our screening and sorting experiments.

**Figure 3.**
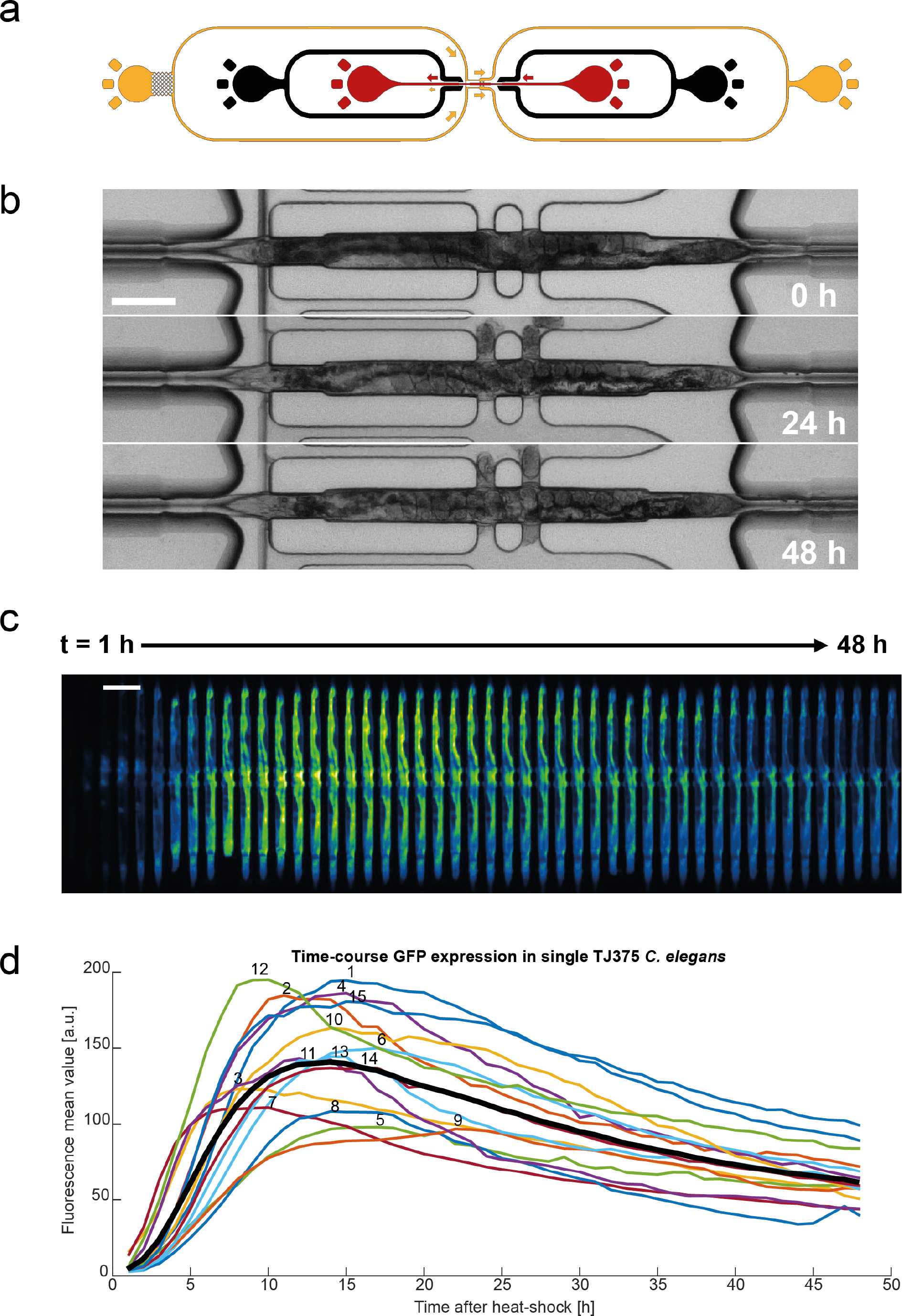
*C. elegans* experimental HSR characterization at individual level. (a) Schematic of the microfluidic design that allows for adult *C. elegans* immobilization. Arrows represent the flow direction. Control layer is represented in black. (b) Images of an immobilized animal at determined time points after trapping. Scale bar represents 100 μm. (c) Increasing time-lapse fluorescence images from an immobilized day-2 adult TJ375 *C. elegans*, after 1-hour heat-shock induction, in microfluidic device along 48 hours, where *t* = 1 h is at the furthest left and *t* = 48 h is at the furthest right. Bar scale = 200 μm. (d) Measured GFP mean expression over time (*t* = 48 h) in individual TJ375 *C. elegans*. Each curve represents a single *C. elegans* (*n* = 15) and they are identified with a number next to the curve. The mean response of the dataset is shown in a bold black line.

### Mathematical modelling identifies translation and degradation rate constants as primary driver of individual variability of dynamics of HSR

Next, we employed a mathematical model of HSR to shed light on the underlying biological mechanism(s) driving the observed heterogeneity among isogenic *C. elegans*. The molecular basis of HSR regulation has been the subject of intense scrutiny in the past, and numerous mathematical models have been proposed to simulate HSR dynamics in several model organisms [33–42]. Here, we adapted an ordinary differential equation (ODE) model of HSR originally developed for human HeLa cells by Scheff *et al.* [43]. This model was among the most comprehensive models of HSR, built on previous modelling works by Petre *et al.*[44] and Szymanska and Zylicz [45]. Briefly, the HSR model describes the network of molecular reactions that responds to an increase in the level of protein denaturation caused by heat insults. The original model parameters were obtained by fitting to literature compiled time-course HSP data of HeLa cells. For the specific purpose of our study and also for a direct comparison to our data of time-course GFP fluorescence in TJ375 (P*hsp-16.2*∷GFP) *C. elegans*, we added a balance equation for the GFP reporter to the original model (details in Methods, Table 1, Table 2 and Figure 4a). Here, we assumed that the rate constants of the GFP transcription and translation mirror those of the HSP, based on the fact that they are controlled by the same promoter. We also set the degradation rate constant of the GFP to equal that of the HSP, since both proteins followed similar dynamics after their peaks in our western blotting experiments (Figure 1d and Supplementary Figure 1).

**Table 1.** Molecular species in the HSR model

| Species | Description |
| --- | --- |
| HSF-1 | Heat shock factor 1 |
| HSF-1<sub>2</sub> | HSF-1 dimer |
| HSF-1<sub>3</sub> | HSF-1 trimer |
| HSE | Heat shock element |
| HSF<sub>3</sub>:HSE | HSF trimer bound to HSE |
| HSP | Heat shock protein |
| HSP:HSF-1 | HSP bound to HSF-1 |
| mRNA | Messenger RNA of HSP and GFP |
| *Prot* | Functional proteins |
| MFP | Misfolded proteins |
| HSP:MFP | HSP bound to MFP |
| GFP | Green fluorescent protein |

**Table 2.**
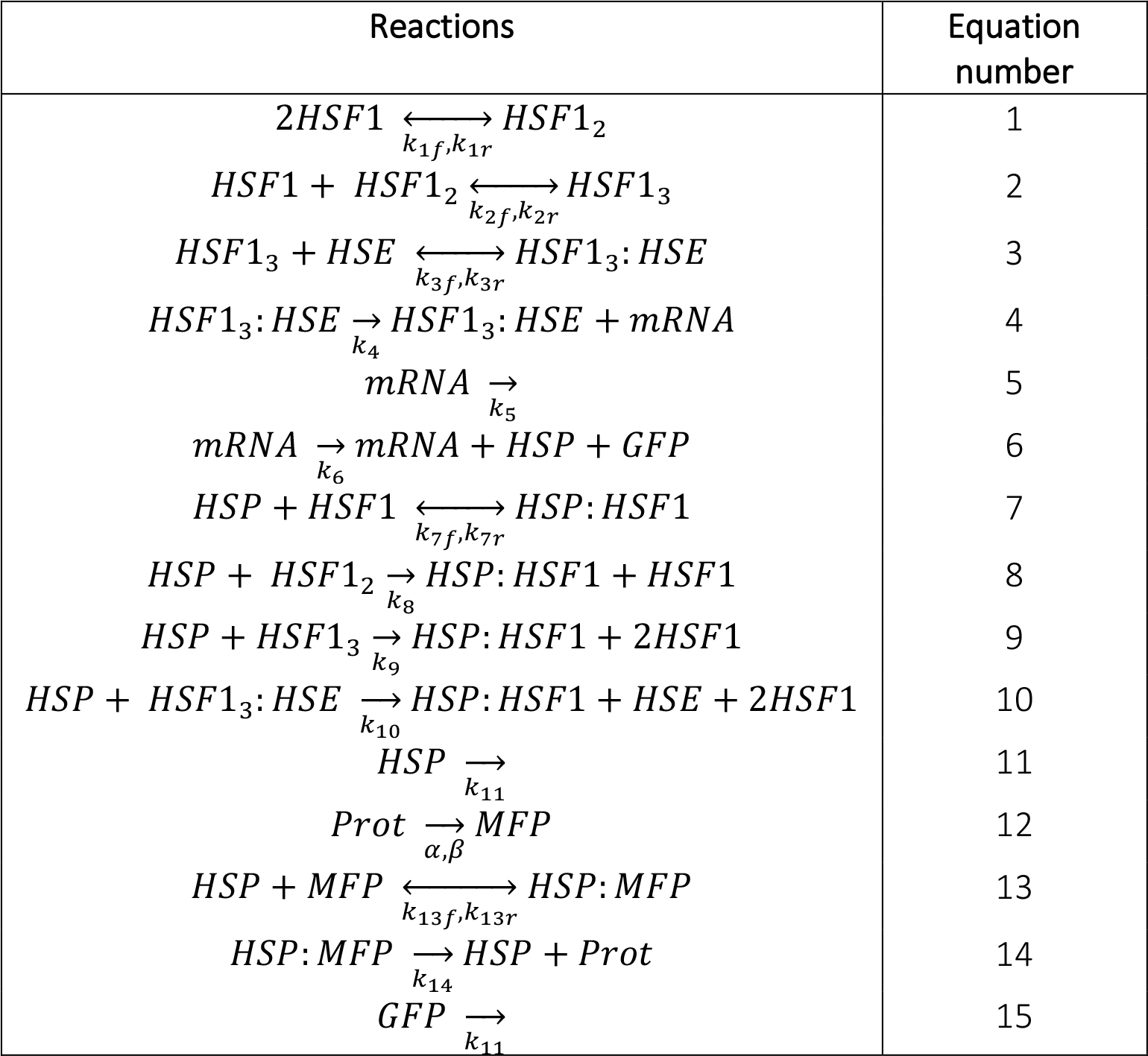
List of reactions in the HSR model

**Figure 4.**
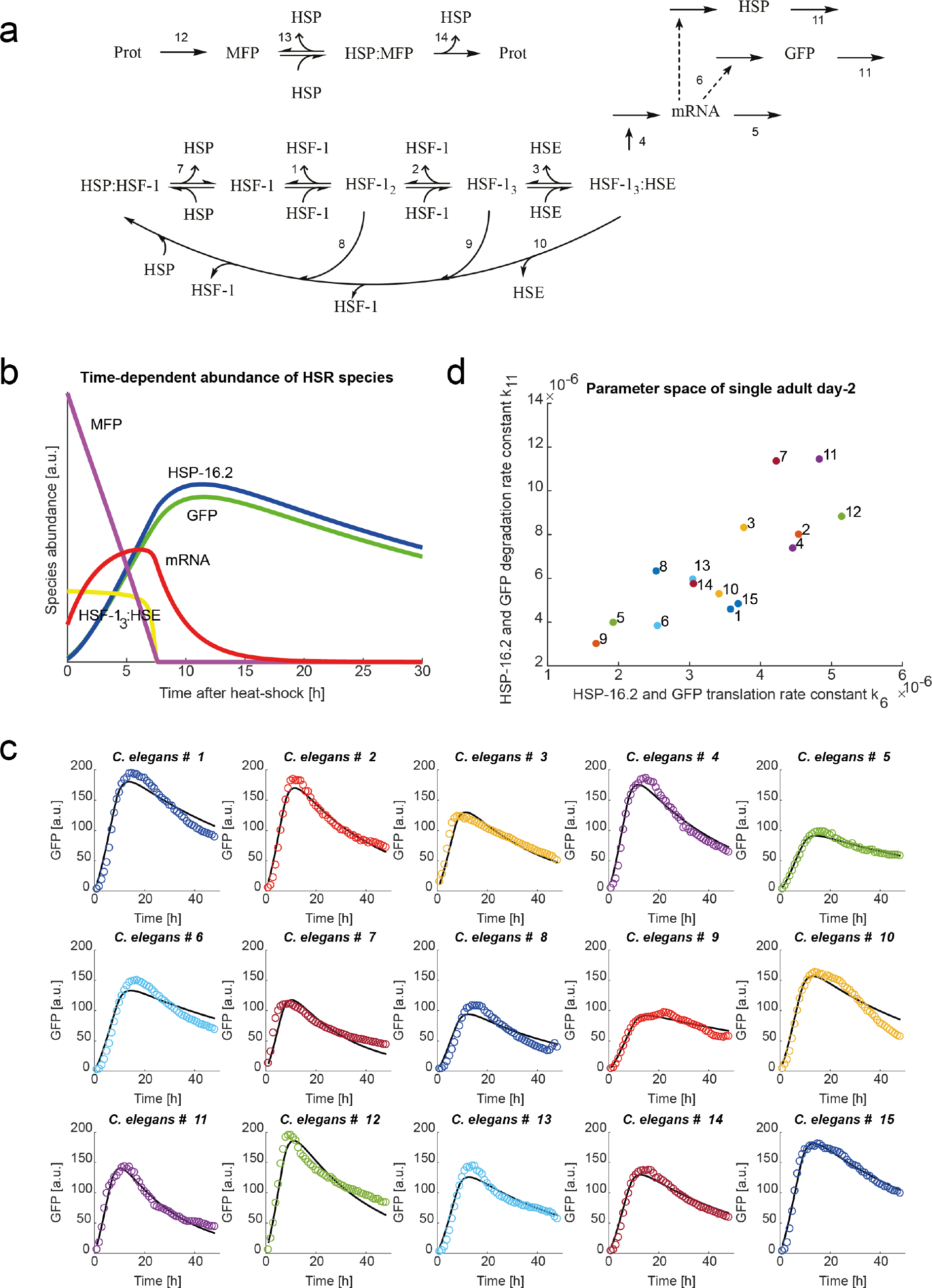
*C. elegans in silico* HSR mathematical modelling. (a) Reaction network describing the *C. elegans* HSR in mutants with GFP as protein reporter of HSP. Solid arrows indicate reaction paths, whereas dotted arrows indicate an activation where the substrate is not consumed. The number annotations correspond to the listed reactions in Table 2. (b) Dynamic expression of the primary species determining HSR: Misfolded proteins (MFP), Trimer HSF-1 bound to HSE (HSF_3_:HSE), mRNA, GFP and HSP-16.2. Heat-shock takes place between −1 h and 0 h. Positive values on the time axis indicate the time in hours after heat-shock. (c) Individual fitting of individual GFP expression for each analysed animal, by varying only parameters *k*_*6*_ and *k*_*11*_ from the average response. Here the solid lines represent the modelled HSR while the circles represent the experimental data. (d) *k*_*6*_ – *k*_*11*_ parameter space found for the experimental dataset shown in Figure 3d. The number next to each data point identifies the *C. elegans* analysed.

We fit the model parameters to the GFP time-course data of individual *C. elegans* in two stages. In the first stage, we fit the GFP concentration predicted by the model to the average *C. elegans* whole-body fluorescence measurements of the fifteen day-2 adult TJ375 *C. elegans* (Figure 3d and Table 3 in Methods section). Figure 4b illustrates the model simulation of the key components of HSR in *C. elegans* after a 1-hour 37° C heat-shock (y-axis not to scale). Misfolded protein (MFP) increases sharply during the 1-hour heat shock (not shown). HSP captures MFP for refolding and MFP levels drop immediately after heat shock. The binding of HSP to MFP frees HSF-1 that would have otherwise formed a complex with HSP. The free HSF-1 undergoes stepwise polymerizations to produce HSF-1 trimers, and after translocating to the nucleus, the trimer binds to the heat-shock element (HSE) to upregulate the expression of HSP (and GFP). As a result, the level of HSP mRNA rises and peaks immediately after MFP returns to its pre-heat shock level. The amount of HSP increases along with its mRNA, though with a slight delay, peaking a few hours later. The GFP signal follows a time-course similar to that of the HSP, consistent with western blotting data (Figure 1c,d).

**Table 3.**
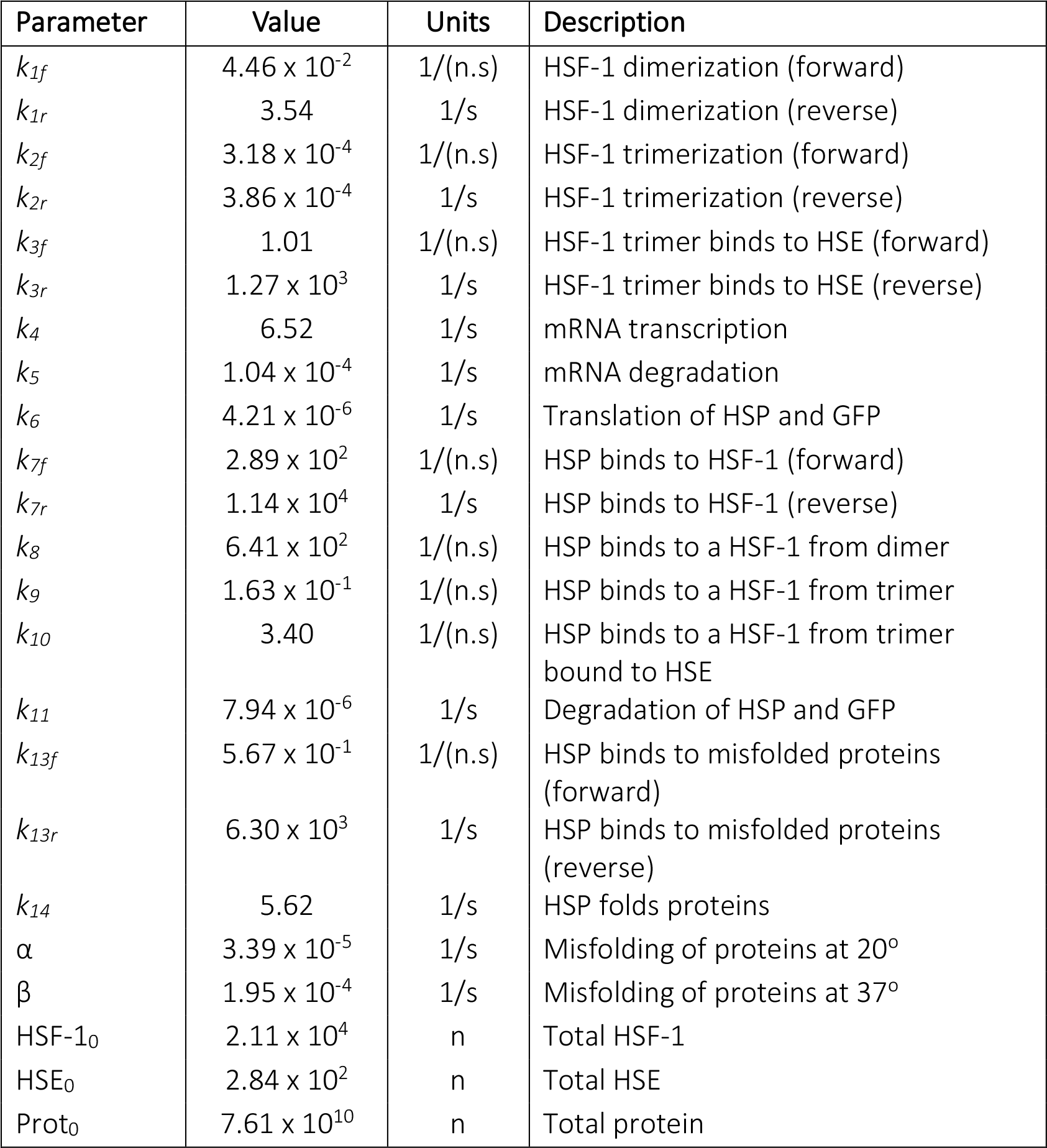
Model parameters estimated for average adult GFP expression data

| Parameter | Value | Units | Description |
| --- | --- | --- | --- |
| $k_{1f}$ | $4.46 \times 10^{-2}$ | 1/(n.s) | HSF-1 dimerization (forward) |
| $k_{1r}$ | 3.54 | 1/s | HSF-1 dimerization (reverse) |
| $k_{2f}$ | $3.18 \times 10^{-4}$ | 1/(n.s) | HSF-1 trimerization (forward) |
| $k_{2r}$ | $3.86 \times 10^{-4}$ | 1/s | HSF-1 trimerization (reverse) |
| $k_{3f}$ | 1.01 | 1/(n.s) | HSF-1 trimer binds to HSE (forward) |
| $k_{3r}$ | $1.27 \times 10^3$ | 1/s | HSF-1 trimer binds to HSE (reverse) |
| $k_4$ | 6.52 | 1/s | mRNA transcription |
| $k_5$ | $1.04 \times 10^{-4}$ | 1/s | mRNA degradation |
| $k_6$ | $4.21 \times 10^{-6}$ | 1/s | Translation of HSP and GFP |
| $k_{7f}$ | $2.89 \times 10^2$ | 1/(n.s) | HSP binds to HSF-1 (forward) |
| $k_{7r}$ | $1.14 \times 10^4$ | 1/s | HSP binds to HSF-1 (reverse) |
| $k_8$ | $6.41 \times 10^2$ | 1/(n.s) | HSP binds to a HSF-1 from dimer |
| $k_9$ | $1.63 \times 10^{-1}$ | 1/(n.s) | HSP binds to a HSF-1 from trimer |
| $k_{10}$ | 3.40 | 1/(n.s) | HSP binds to a HSF-1 from trimer bound to HSE |
| $k_{11}$ | $7.94 \times 10^{-6}$ | 1/s | Degradation of HSP and GFP |
| $k_{13f}$ | $5.67 \times 10^{-1}$ | 1/(n.s) | HSP binds to misfolded proteins (forward) |
| $k_{13r}$ | $6.30 \times 10^3$ | 1/s | HSP binds to misfolded proteins (reverse) |
| $k_{14}$ | 5.62 | 1/s | HSP folds proteins |
| $\alpha$ | $3.39 \times 10^{-5}$ | 1/s | Misfolding of proteins at 20° |
| $\beta$ | $1.95 \times 10^{-4}$ | 1/s | Misfolding of proteins at 37° |
| HSF-1<sub>0</sub> | $2.11 \times 10^4$ | n | Total HSF-1 |
| HSE<sub>0</sub> | $2.84 \times 10^2$ | n | Total HSE |
| Prot<sub>0</sub> | $7.61 \times 10^{10}$ | n | Total protein |

In the second stage, we performed parameter fitting to the GFP time-courses of individual *C. elegans* and obtained 15 different parameter sets, one for each *C. elegans* in the experiment. Here, we assumed that the parameter values of individual animals differ amongst each other and from the average *C. elegans* only for a small number of parameters. More specifically, we set the majority of the model parameters to the values obtained from the fitting of the average whole-body GFP fluorescence above, and re-fit only a select set of parameters to the single-animal whole-body GFP data. To select the set of parameters that are able to fit the single-animal GFP time-course, we performed a parametric sensitivity analysis and considered parameters that are related to biologically regulated processes (such as transcription, translation and degradation rate constants) (details in Methods). Guided by the result of the sensitivity analysis (Supplementary Figure 6 and Methods), we selected two parameters related to the protein turnover, specifically the protein translation and degradation rate constants. Indeed, using only these two parameters (see Methods for detail), we were able to obtain accurate model fits to the GFP time-courses of individual *C. elegans* (Figure 4c). The distribution of the protein turnover parameters for the individual *C. elegans* is shown in Figure 4d, where the numbers next to the parameter points correspond to the identity of the single *C. elegans* in Figure 3d. The results suggest that differences in the protein turnover capacity are a major driver of the heterogeneity in the HSR dynamics among individual *C. elegans* in isogenic population.

### The abrupt collapse of proteostasis during early adulthood contributes to the wide-spread of heterogeneous responses to heat-shock

Protein translation, trafficking and degradation are essential for the functionality of cellular processes. Upon a proteotoxic stress, such as heat-shock, the cell upregulates molecular chaperones, including HSP, to help re-establish protein homeostasis or proteostasis [46–49]. A natural decline in the activity of HSR and in survival to thermal stress in *C. elegans* coincides with a proteostasis collapse in multiple tissues, starting as early as 12 hours into adulthood (day-1 adults) [50–52]. In the above experiments, we used day-2 adult *C. elegans* that may have potentially already experienced such proteostatic decline. We thus hypothesized that younger *C. elegans* should demonstrate a “better” response to the same heat-shock stress. To confirm this hypothesis, we repeated our assessments of HSR dynamics for a cohort of day-1 adults exposed to the same 1-hour 37° C heat-shock treatment.

The HSR assessment using our high-throughput microfluidic system indicated that the average whole-body GFP expression of day-1 adult TJ375 population reaches its maximum earlier than day-2 adults (8 hr vs. 12 hr after heat shock; Figure 5a and 5b). In order to further confirm that the source of the differences between these two cohorts is associated with proteostatic decline, we characterized the single-animal HSR dynamics of day-1 adult *C. elegans* (*n* = 10) using time-lapse fluorescence microscopy (Figure 5c, Supplementary Figure 7a). We found that the GFP time courses of day-1 adults have, on average, a higher maximum fluorescence value than day-2 adult nematodes (195.11 ± 63.78 a.u. for day-1 vs 147.74 ± 35.16 a.u. for day-2 adults), and also that they peak earlier (8.7 ± 1.5 h for day-1 vs 14 ± 3.48 h for day-2). We then applied our mathematical model to the combined GFP expression time course curves from day-1 and day-2 TJ375 adults (*n* = 25) and repeated the two-stage parameter fitting process (Supplementary Figure 7b). Again, by varying translation and degradation rate constants among individual nematodes, we were able to explain the variability of HSR dynamics among individual nematodes.

**Figure 5.**
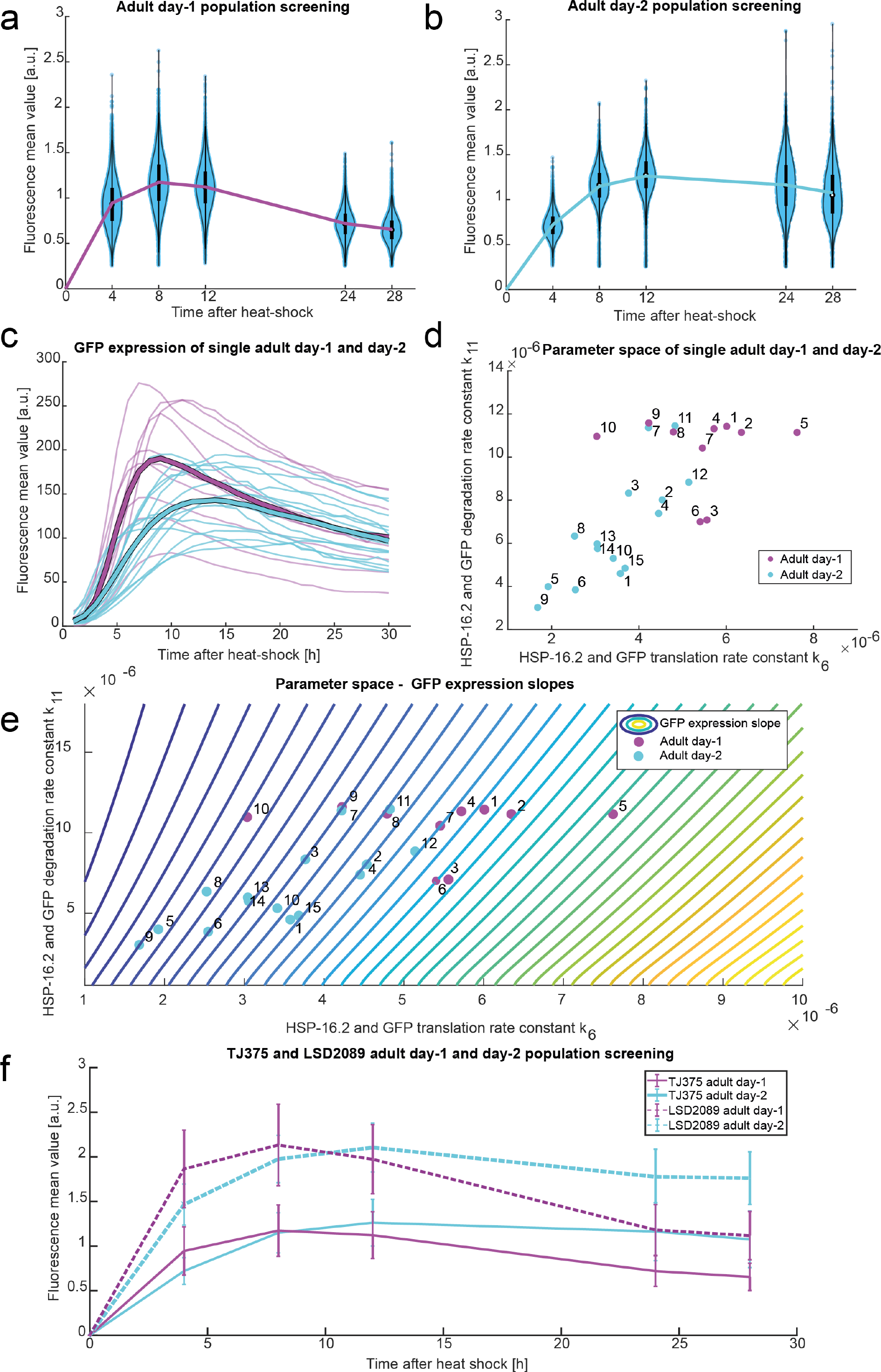
HSR study for *C. elegans* with different ages (day-1 vs day-2 adults). (a) Violin plots of the fluorescence mean values of individual *C. elegans* measured in an isogenic cohort at day-1 adulthood (*n* = 6917) at determined time points (4, 8, 12, 24 and 28 hours) after heat-shock. The change in the mean value of each group at each sorting or screening time is shown with the magenta line. (b) Homologous violin plots of the fluorescent mean values of individual *C. elegans* measured in an isogenic cohort at day-2 adulthood (*n* = 2716). (c) GFP mean expression in single animals over time (*t* = 30 h): In magenta day-1 adult *C. elegans* (*n* = 10) and in cyan day-2 adult *C. elegans* (*n* = 15). (d) *k*_*6*_ – *k*_*11*_ parameter space found for the experimental datasets shown in (c). The number next to each data point identifies the *C. elegans* analysed (See Figure 3d and Supplementary Figure 7), where magenta dots correspond to day-1 adults and cyan dots to day-2 adults. (e) Contour plot of GFP expression slope values according to *k*_*6*_ and *k*_*11*_ values (translation and degradation rates of GFP), where dark blue represents lower values and yellow higher values. Steeper slope isolines correlate with higher translation rates and lower degradation rates. Overlaid are the measured parameters shown in (d). (f) GFP expression curves of LSD2089 (*ncl-1*, P*hsp-16.2*∷GFP) (dotted lines) and TJ375 (P*hsp-16.2*∷GFP) (solid lines) for adult day-1 (in magenta) and adult day-2 (in cyan) after heat-shock. In each population, for each time point we show the mean fluorescence value and its standard deviation, when measured with our high-throughput microfluidic system.

The distribution of the protein translation and degradation rate constants for day-1 and day 2 adult *C. elegans* are depicted in Figure 5d (Supplementary Table 2). The results of the model fitting show that younger *C. elegans* (day-1 adults) have on average higher rate constants for protein translation and degradation, both of which are indicators of a higher proteostatic capacity, confirming our hypothesis. Besides the lower protein turnover parameters, the day-2 cohort also displays a higher heterogeneity (variance) in the rate constants. The increased heterogeneity agrees with a previous study of protein turnover in *C. elegans* ageing [53]. Moreover, when comparing the GFP expression curves of day-1 and day-2 adult *C. elegans* (Figure 5c), we observed an overlap, where a fraction of day-2 adult *C. elegans* (especially, day-2 *C. elegans* number 7, 11 and 12) exhibited GFP expression curves that are similar to the typical GFP expression curves of day-1 adult *C. elegans*. A further comparison of the protein turnover parameters shows that these specific day-2 adult *C. elegans* indeed have translation and degradation rate constants that are higher than those of the other day-2 animals and closer to those of day-1 adults (Figure 5d and Supplementary Table 2). These comparisons suggest that ageing-associated decline of protein turnover does not occur at equal rates for nematodes in a given population. Indeed, other studies have reported a significant stochasticity in the onset and the rate of deterioration in protein turnover within isogenic *C. elegans* populations [54].

We then investigated further the impact of the protein turnover heterogeneity on HSR capacity. For this purpose, we focused on the early time-course of HSR as an indicator of its capacity. More specifically, we associated the HSR capacity in individual *C. elegans* with the speed and strength of the GFP expression increase in response to a heat-shock. We used the HSR model to simulate the time to reach the maximum GFP expression – a marker of speed – and the maximum level of the GFP expression – a marker of strength of the HSR. Model simulations show that the speed of the HSR in individual *C. elegans* is positively correlated with their protein translation rate constants (Supplementary Figure 7c), *i.e.* a higher translation rate constant gives a faster GFP increase in response to heat-shock. Meanwhile, the maximum GFP expression level is proportional to the ratio between protein translation and degradation rate constants among single *C. elegans* (Supplementary Figure 7d).

We subsequently computed the slope of GFP expression by taking the ratio between the maximum GFP level and the time to reach the maximum and used this value as a single combined marker for HSR capacity in individual *C. elegans*. Figure 5e depicts the isolines (contours) of the GFP slopes as a function of the rate constant of protein translation (x-axis) and protein degradation (y-axis) (details in Methods). The isolines show that a faster translation corresponds to a higher slope, whilst a higher degradation corresponds to a lower slope. Thus, the effect of declining protein translation on the HSR capacity can be abrogated by lowering protein degradation. Furthermore, the protein translation rate constant exerts a stronger influence on the GFP slope than the protein degradation rate constant. But, the dependence of the slope on protein translation weakens with increasing level of protein degradation, as indicated by denser isolines at lower degradation rate constants. By comparing the locations of the parameters of day-1 adult *C. elegans* with those of day-2 adult *C. elegans* with respect to the isolines on Figure 5e, we noted that day-1 adult *C. elegans* have generally higher GFP slope than day-2 adults (see Supplementary Table 3, two-sample *t* test p-value < 0.0001). Thus, the decline in the protein turnover during early adulthood is detrimental to the HSR capacity of the animal.

In order to confirm the influence of the protein translation rate constant on the HSR capacity, we crossed knocked down *ncl-1(e1942)* mutants that have increased amounts of ribosomes and almost double the amount of general protein translation [55], into TJ375 (P*hsp-16.2*∷GFP), producing a transgenic strain LSD2089 (P*hsp-16.2*∷GFP, *ncl-1(e1942)*). We measured the GFP expression time-course of heat shocked day-1 and day-2 adults LSD2089 *C. elegans* and compared them with that of TJ375 (Figure 5f). We found that while the times to reach the maximum GFP expression in the populations are similar between same age LSD2089 and TJ375, the mean GFP expression values of LSD2089 are 1.8 times (*e.g.*, almost doubled) compared TJ375. The higher GFP slopes in the crossed strain is consistent with the trend predicted by the isolines in Figure 5e – a higher protein translation rate constant corresponds to a higher slope. The deviation in the GFP time-courses between day-1 and day-2 adults of LSD2089 *C. elegans* mirrors that of TJ375. These data thus suggest that despite an improved HSR activity conferred by a higher protein translational capacity, LSD2089 *C. elegans* still experience a similar proteostatic collapse during its early adulthood.

### Mathematical modelling to identify the early predictive features of the HSR on *C. elegans* lifespan

Previous studies showed that the strength of HSR in TJ375 *C. elegans* as measured by their HSP expression levels is predictive of lifespan [20, 28]. Based on our modelling results, which point out that assayed *C. elegans* of the same chronological age appear to have different protein-turnover capacities, we hypothesize that the HSR capacity, and thus the protein-turnover machinery of each animal, is the underlying element of this lifespan predictor. For hypothesis validation, we performed model simulations to determine the sorting criterion that demarcates the top and bottom HSR capacity among day-2 adult *C. elegans.* Briefly, we generated and simulated an ensemble of 10 000 *in-silico C. elegans* models, where the models differed among each other only in the values of protein translation and degradation rate constants (Figure 6a,b). The model parameters were generated by a random sampling of the parameters, assuming a bivariate Gaussian distribution with the mean and variance equalling to those of single *C. elegans* parameters in Figure 5c. We simulated each single-animal model to produce a GFP expression time-course. The collection of 10 000 of such time courses is depicted in Figure 6b. The simulated GFP curves agree well with our screening measurements from the high-throughput experiments, as shown in Figure 1g (Supplementary Figure 8), as well as with the sorting-screening experiment shown in Figure 2c (Supplementary Figure 9).

**Figure 6.**
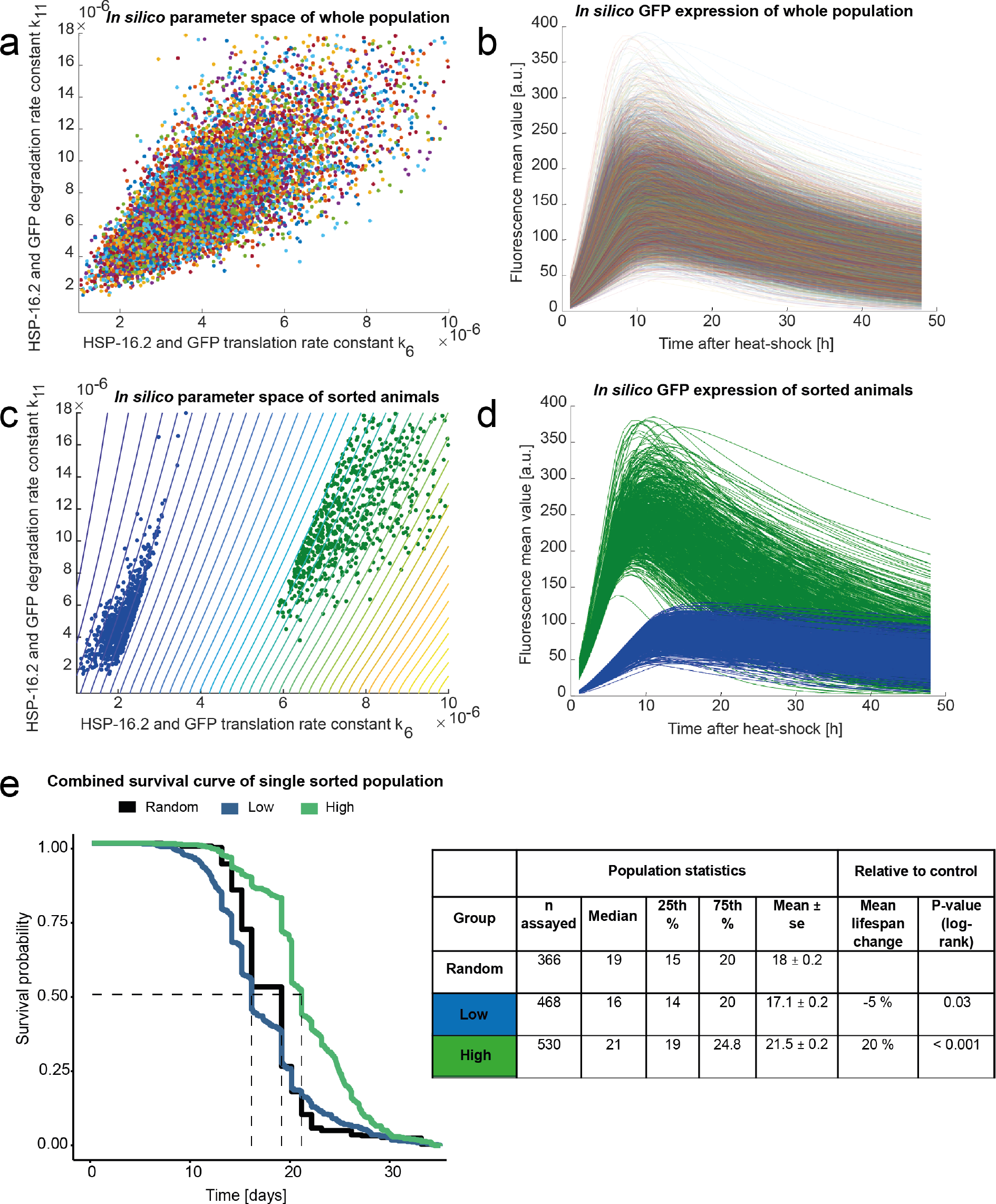
Sorting and lifespan studies based on HSR difference. (a) *In silico k*_*6*_ – *k*_*11*_ parameter space (*n* = 10 000) created around the experimental data from the individual HSR characterization (*n* = 25) shown in Figure 5d. (b) Simulated profiles (*n* = 10 000) of GFP expression after heat-shock, where each curve is generated with a unique combination of *k*_*6*_ and *k*_*11*_, from (a). (c) Parameter subspaces for groups ‘high’ and ‘low’ which correspond to GFP expression curves with different slopes, when selecting top and bottom 5% of the GFP expression, 5 hours after heat-shock. Overlaid it is shown the contour plot of Figure 5e. (d) Selected GFP expression profiles with the sorting criteria top and bottom 5% GFP expression at 5 h after heat-shock. (e) Lifespan curves resulting from the lifespan assays of the sorted *C. elegans* groups (combined results from 3 biological repeats). The population statistics are enlisted on the right.

Based on the simulations of the model ensemble in Figure 6a, we found that by taking the top and bottom 5% GFP expression of the population at 5 hours post heat-shock will results in significantly different HSR capacity with the average GFP slope of the top 5% being 4-fold higher than that of the bottom 5% (25.54 a.u./h for “high” group vs 6.41 a.u./h for “low” group). The sorting criteria above would capture *C. elegans* corresponding to the parameter space depicted in Figure 6c (the top 5% in green and the bottom 5% in blue) and to the GFP expression time-courses shown in Figure 6d. The model simulations shown in Figure 6d suggest that the two sub-populations should comprise *C. elegans* with starkly different GFP slopes and thus HSR capacities. We implemented the sorting criterion as determined by the above model simulations in a sorting experiment using a day-2 adult TJ375 *C. elegans* population following heat-shock. We used our high-throughput sorting microfluidic device (Figure 2a) to sort and isolate the two sub-populations. Figure 6d gives the combined survival curves for the top and bottom 5 % GFP expressing day-2 adult *C. elegans* at *t* = 5 h after heat-shock for 3 biological repeats; 2 of those repeats were obtained using a fully-automated lifespan machine [56] and one by manual inspection, as detailed in Methods and Supplementary Data File 1. In combination, the top 5% sub-population has a 4.4-day (25%) higher mean lifespan than the bottom 5% in these independent experiments (p-value < 0.001 Figure 6e, Supplementary Data File 1b-d). The result of our lifespan study using model-based sorting criteria thus validates our hypothesis that the HSR capacity – as measured by the GFP slope – is an early biomarker for longevity among a genetically identical *C. elegans* population – as early as adult day-1.

### Heterogeneity of HSR time-course is visible at the tissue level

Individual tissues within an organism have been shown to respond to not only their own proteostasis disruption but also that of adjacent tissues in a coordinated fashion [57]. Tissue-specific HSR regulation has further been tied to distinct cellular proteomes and specific profiles of heat-shock inducible genes, which give particular tissue-dependant contributions to the whole-organismal proteostasis machinery [58]. To examine tissue-specific HSR dynamics in *C. elegans*, we reanalysed the above time-lapse images of individual day-1 and day-2 adult nematodes upon heat-shock. More specifically, we sectioned the images into five equal-sized compartments and monitored the temporal expression of GFP in distinct tissue regions, namely head, anterior intestine, central intestine, posterior intestine, and tail (Figure 7a). In each of these tissue sections, we obtained the mean GFP expression over 30-hour time course for day-1 adult (Figure 7b) and day-2 adult *C. elegans* (Figure 7c). Figure 7d (Supplementary Table 4) compares the GFP expression of the day-1 and day-2 adult datasets in different tissue sections.

Comparison across compartments indicated that the intestine central compartment exhibits the quickest response to heat shock, followed by its neighbouring sections and then the extremities, *i.e.* the head and tail. The head section of both ages has its maximum GFP expression value at the approximately the same time (9.7 h vs 10.3 h after heat-shock for day-1 and day-2, respectively). But, in other compartments the GFP expression of the day-1 cohort has a faster rise than the day-2 population. In the intestine anterior, the time to reach the maximum GFP is 10 h for day-1 versus 13.5 h for day-2 (Figure 7b-d). In the intestine central section, day-1 nematodes take 8.4 h on average while day-2 animals take 12.6 h on average to reach the GFP peak (Figure 7b-d). In the intestine posterior compartment, the peak times are 9 h for day-1 and 16.7 h for day-2 (Figure 7b-d). Finally, in the tail section, the time to achieve maximum GFP expression is 10.1 h for day-1 and 16.7 h for day-2. Besides faster times to reach peak GFP, day-1 nematodes also have a higher average maximum expression across most of the compartments, except for the tail. Taken together, the comparison above indicates that the early adulthood decline of HSR in the intestine is notably stronger than in other tissues. This suggests that the proteostasis collapse [50, 51] of the intestine may underlie the heterogeneity in older individuals – as old as adult day-2.

**Figure 7.**
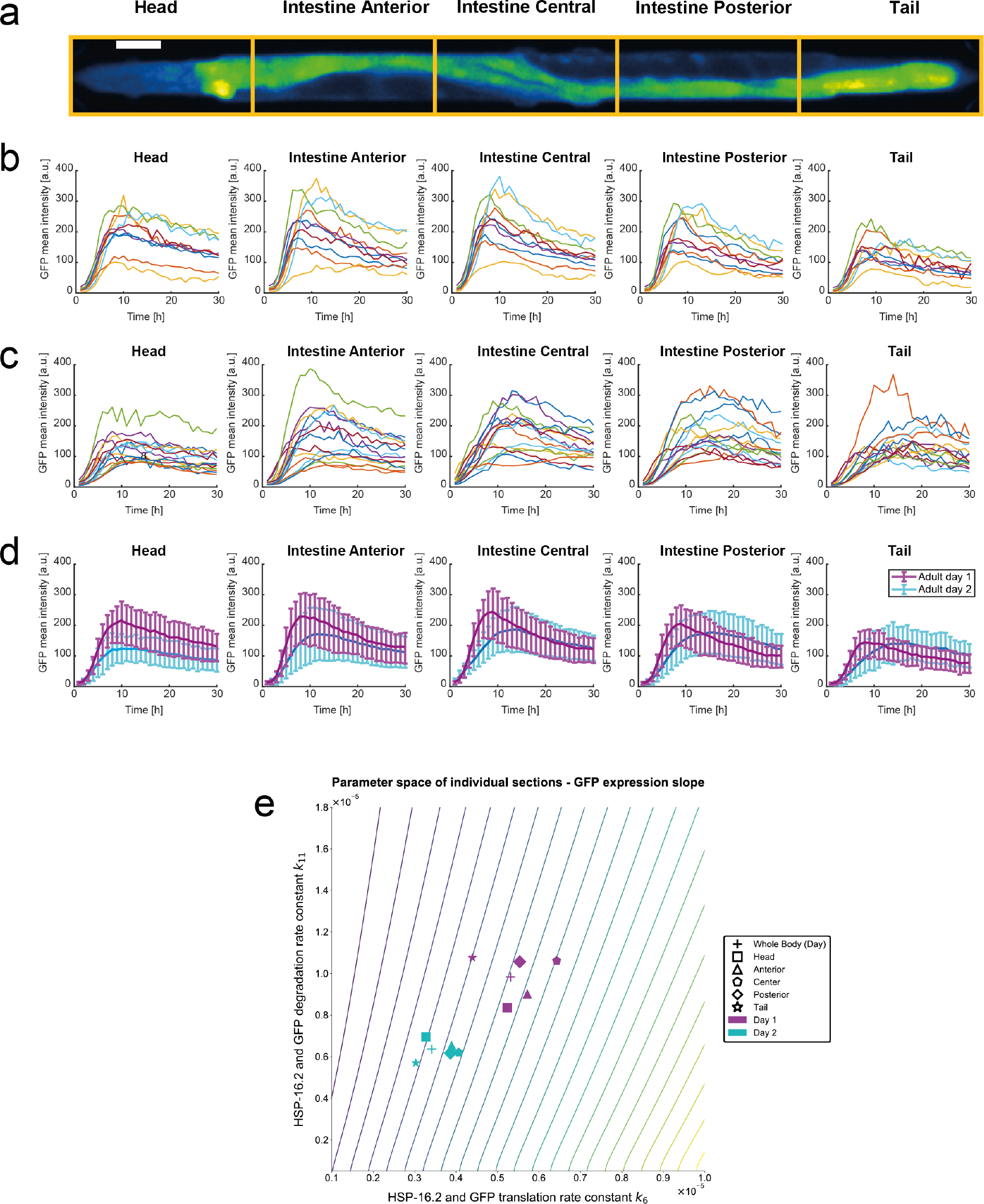
Differences in tissue-specific HSR in *C. elegans*. (a) Fluorescence image of an immobilized day-2 adult TJ375 *C. elegans* in a microfluidic device for long-term immobilization; we indicated with yellow boxes the segmentation for tissue GFP expression analysis, namely head, intestine anterior, intestine central, intestine posterior and tail. Bar scale = 50 μm. (b) GFP mean expression over time (*t* = 30 h) of different sections of the body of adult day-1 *C. elegans* after heat-shock. (c) GFP mean expression over time (*t* = 30 h) of different sections of the body of adult day-2 *C. elegans* after heat-shock. (d) Average GFP mean expression over time (*t* = 30 h) for day-1 (in magenta) and day-2 (in cyan) adult *C. elegans* of different sections of the body. Error lines represent the standard deviation of the dataset. (e) Parameter space of individual sections of *C. elegans* adult day-1 (in magenta) and day-2 (in cyan). Overlaid are the isolines of GFP expression slope shown in Figure 5e.

We also used the mathematical model above to describe the compartment-specific HSR dynamics, following the same parameter estimation procedure aforementioned. As before, we found that the heterogeneity of GFP expression dynamics across the compartments can be explained by varying the model parameters associated with protein translation and degradation. Figure 7e depicts the rate constants of protein translation and degradation estimated using the average GFP dynamics of each compartment for day-1 and day-2 nematodes (see Supplementary Figure 10 for the full parameter estimation of individual compartments in single *C. elegans*). Comparing across compartments, the intestine central section shows the highest GFP slope, followed by the intestine anterior, the intestine posterior, the head, and lastly the tail section. Comparison of day-1 and day-2 adults confirmed the aforementioned trend for each compartment; specifically that there is an age-related decline in the protein translation and degradation (protein turnover) and consequently in the GFP slope and therefore the HSR capacity (Figure 7e). The decline in the GFP slope again differs across compartments with those toward the middle of the animal exhibiting a larger drop. Notably, in comparison to day-1 individuals, the discrepancies of the GFP slopes among the compartments become less pronounced for day-2 cohort. In other words, the HSR capacity becomes more uniform across the compartments from day-1 to day-2, which is opposite to the trend for animal-to-animal heterogeneity. That said, within each compartment, we still observed an increase in the heterogeneity of protein turnover parameters with age (Supplementary Table 5).

### Maternal contribution to HSR heterogeneity is associated with protein translational capacity of offspring embryos

Given the above observation of proteostatic decline with age in adult nematodes and its effects on HSR capacity, we reasoned that proteostasis and HSR capacity in embryos should be reset to optimal efficiency and individual embryos’ HSR should be similar to each other. To test this hypothesis, we quantified the GFP expression of embryos from TJ375 (P*hsp-16.2*∷GFP). Briefly, we submitted gravid day-1, day-2 and day-4 *C. elegans* hermaphrodites to heat-shock for 1 hour at 37° C and immediately after collected their embryos. We placed the embryos in a microfluidic device for a longitudinal observation, as shown in Figure 8a-b (described in the Methods section) and recorded their GFP fluorescence intensities every 10 minutes. The GFP fluorescence time-courses for the embryos are shown in Figure 8c, where the average responses of each dataset are indicated in bolder lines (magenta for embryos of day-1 adults, cyan for embryos of day-2 adults, and green for embryos of day-4 adults).

**Figure 8.**
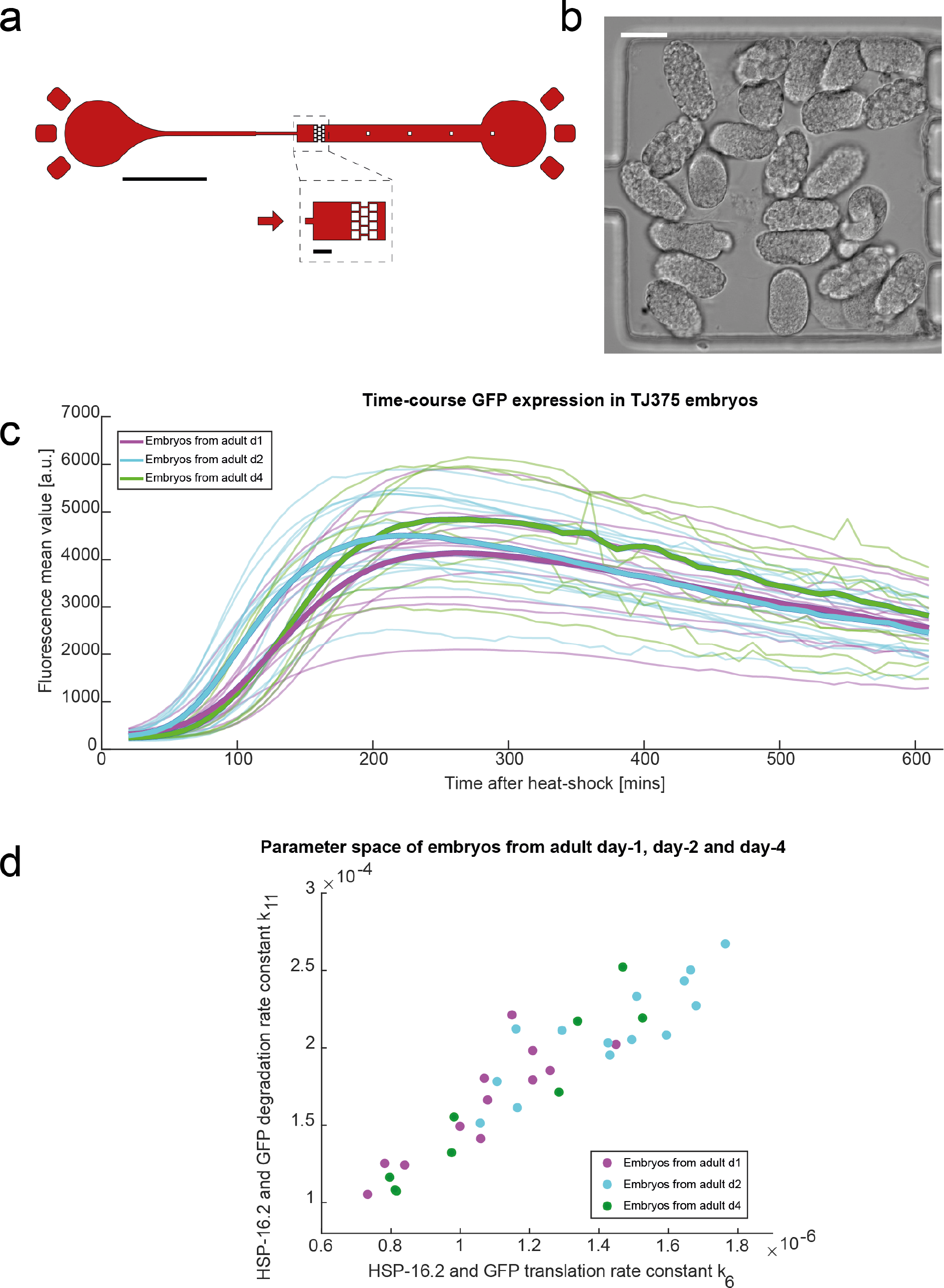
HSR in embryonic *C. elegans*. (a) Schematic of the microfluidic design that allows for long-term embryonic *C. elegans* observation. Arrows represent the flow direction. Bar scales represent 1 mm and 100 μm. (b) Bright field image of a collection of embryos in the microfluidic chamber filled with embryos. Bar scale represents 30 μm. (c) GFP expression after heat-shock of embryos coming from different age gravid *C. elegans*. Magenta lines represent eggs from day-1 adults, cyan lines eggs from day-2 adults and green lines eggs from day-4 adults; over time (*t* = 10 h). Bolder lines represent the average response of each dataset. (d) *k*_*6*_ – *k*_*11*_ parameter space of the embryos shown in (c). Diamonds in magenta represent embryos from adult day-1, in cyan embryos from adult day-2 and in green embryos from adult day-4.

In comparison to adults, HSR dynamics in embryos are significantly faster, with average GFP fluorescence peaking in 5 hours after heat-shock. (Figure 8c). The faster HSR dynamics suggests a higher HSR capacity in embryos than in adults, confirming our hypothesis. That said, there is still heterogeneity in the HSR time-courses among individual embryos (Figure 8c). We applied our mathematical modelling to the embryo dataset, estimating the model parameters that reproduce the average response among the embryos (Table 4) and then finding the protein translation and degradation rate constants for individual embryos (Figure 8d, Supplementary Figure 11, Supplementary Table 7). As for adults, variation of protein translation and degradation rate constants was able to capture the heterogeneity in the HSR dynamics among embryos. In the context of evolutionary trade-off related to developmental speed to reach reproduction, the embryos of day-1 mothers have slower protein turnover rate and thus a lower HSR capacity than those of day-2 mothers (*k*_6_: *p*-value = 4.2×10^−4^, *k*_11_: *p*-value = 2.4×10^−3^, two-sample *t*-test on mean difference). Also, in comparison to embryos of day-4 mothers, embryos of day-2 mothers have faster protein turnover and HSR dynamics (*k*_6_: *p*-value = 9.3×10^−3^, *k*_*11*_: *p*-value = 1.8×10^−2^, two-sample *t*-test on mean difference), suggesting that the HSR capacity in embryos eventually declines with the mothers age [19].

**Table 4.** Model parameters estimated for average embryo GFP expression data

| Parameter | Value | Units | Description |
| --- | --- | --- | --- |
| $k_{1f}$ | $1.39 \times 10^{-3}$ | 1/(n.s) | HSF-1 dimerization (forward) |
| $k_{1r}$ | 7.15 | 1/s | HSF-1 dimerization (reverse) |
| $k_{2f}$ | $8.16 \times 10^{-6}$ | 1/(n.s) | HSF-1 trimerization (forward) |
| $k_{2r}$ | $2.44 \times 10^{-4}$ | 1/s | HSF-1 trimerization (reverse) |
| $k_{3f}$ | 2.28 | 1/(n.s) | HSF-1 trimer binds to HSE (forward) |
| $k_{3r}$ | $7.58 \times 10^3$ | 1/s | HSF-1 trimer binds to HSE (reverse) |
| $k_4$ | $1.68 \times 10$ | 1/s | mRNA transcription |
| $k_5$ | $4.60 \times 10^{-5}$ | 1/s | mRNA degradation |
| $k_6$ | $1.16 \times 10^{-6}$ | 1/s | Translation of HSP and GFP |
| $k_{7f}$ | 6.25 | 1/(n.s) | HSP binds to HSF-1 (forward) |
| $k_{7r}$ | $3.41 \times 10^4$ | 1/s | HSP binds to HSF-1 (reverse) |
| $k_8$ | $8.87 \times 10^2$ | 1/(n.s) | HSP binds to a HSF-1 from dimer |
| $k_9$ | $8.05 \times 10^{-2}$ | 1/(n.s) | HSP binds to a HSF-1 from trimer |
| $k_{10}$ | 8.15 | 1/(n.s) | HSP binds to a HSF-1 from trimer bound to HSE |
| $k_{11}$ | $1.72 \times 10^{-4}$ | 1/s | Degradation of HSP and GFP |
| $k_{13f}$ | $8.73 \times 10^{-1}$ | 1/(n.s) | HSP binds to misfolded proteins (forward) |
| $k_{13r}$ | $1.51 \times 10^2$ | 1/s | HSP binds to misfolded proteins (reverse) |
| $k_{14}$ | $3.75 \times 10^2$ | 1/s | HSP folds proteins |
| $\alpha$ | $6.08 \times 10^{-5}$ | 1/s | Misfolding of proteins at 20° |
| $\beta$ | $2.12 \times 10^{-4}$ | 1/s | Misfolding of proteins at 37° |
| HSF-1<sub>0</sub> | $5.05 \times 10^2$ | n | Total HSF-1 |
| HSE<sub>0</sub> | $4.51 \times 10$ | n | Total HSE |
| Prot<sub>0</sub> | $9.90 \times 10^8$ | n | Total protein |

In terms of evolution, there is a trade-off between developing and reproducing rapidly and the quality of the embryos [19]. For instance, the hatching size, growth rate, developmental speed, starvation resistance and fecundity, all show heterogeneity stemming from the maternal’s age and packaging of the oocytes, *i.e.*, day-1 mothers produce lower quality embryos, day-2 mothers the highest quality embryos, and then the embryo quality declines with the mother’s age [19]. Our modelling revealed a maternal contribution to the heterogeneity of HSR dynamics, in this case corresponding to the mother’s age, similar to these other heterogeneous phenotypes of *C. elegans*. Embryos from day-1 and day-2 mothers had approximately an equal level of heterogeneity in protein turnover rates *k*_6_ and *k*_11_, with day-2 mothers producing the least heterogeneous embryos. Meanwhile, day-4 mothers had the most heterogeneous embryos. Thus, in the context of HSR capacity and its heterogeneity, day-2 mothers produced the optimal embryos. These results suggest that while proteostasis should be reset in embryos, there still exist heterogeneity among the embryos, which may arise from the packaging of nutrients, mRNA, and proteins from the mother to the egg [19].

## Discussion

Lifespan varies greatly within a population, even among genetically identical animals reared in the same environment [13]. While the heterogeneity of heat shock responses (HSR) or other stress responses has been associated with individual life expectancy [8–13], the underlying molecular determinants of such heterogeneity are still poorly understood. Through the use of novel microfluidic system, which enable high-throughput and high resolution HSR experiments, and molecular level mathematical modelling of the HSR, we identified that:

1. The HSR of individual *C. elegans* follows different HSR dynamics;
2. The heterogeneity of HSR dynamics are contributed by differences in the protein translation and degradation (*i.e.* protein turnover) among single *C. elegans*;
3. There is a rapid decline in the protein turnover (proteostatic collapse) in early life – as early as day-1;
4. The amount of age-related decline in protein turnover is not uniform in an isogenic *C. elegans* population, which results in an increase in the protein turnover variability among the animals;
5. The decline in protein turnover leads to a lowered HSR activity;
6. The decline of protein translation and degradation during early adulthood results in the phenotypic variance of lifespans of these isogenic *C. elegans* reared in the same environment. Specifically, nematodes having higher HSR capacity demonstrate longer mean lifespan;
7. The heterogeneity of HSR dynamics is also seen within compartments of individual nematodes with compartments toward the middle of the animal having a higher HSR capacity;
8. The HSR capacity of each compartment also declines with age, but this decrease varies across the compartment with the central intestine compartment showing the largest drop;
9. As a result, the HSR capacity become more uniform from day-1 to day-2
10. Nevertheless, the animal-to-animal variability of protein turnover within each compartment increases with age, again signifying the contribution of uneven proteostatic decline to the heterogeneity in the HSR among the nematodes.

We found that the response to heat-shock as measured by GFP expression from a heat shock promoter is remarkably heterogeneous, yet at the same time, reproducible. By longitudinal re-sampling of a heat-shocked population, we found no change in noise as defined by the standard deviation divided the mean [59, 60] for between 4 and 48 hours after heat shock (Figure 1g, Supplementary Figure 3). In eukaryotes, random fluctuations in chromatin states or in the number of ribosomes can be a major source of variation in gene expression [59, 60]. However, altering chromatin remodeller which are important for the HSR (*i.e.*, *jmjd-3.1* mutants [32]; Supplementary Figure 4) or mutant backgrounds with higher numbers of ribosomes (*i.e., ncl-1* mutants; Figure 5f) did not affect the spread in standard deviation nor the overall heterogeneity.

To identify the molecular underpinning or major driver of the observed heterogeneity in HSR activity, we applied mathematical modelling to our high-resolution population and individual HSR GFP fluorescent measurements. We discovered that we can explain our experimental data of HSR time-courses in adult day-1 and day-2 nematodes by adjusting only two model parameters: protein translation rate constant *k*_*6*_ and protein degradation parameter *k*_*11*_ (Figure 4). We observed a strong link between these two parameters, whereby animals with a higher translation rate also showed a larger degradation rate, and vice versa (Figure 5). Furthermore, we observed a rapid age-dependent decline in the activity of the HSR between day 1 and day 2 adults that strongly influenced the two protein turnover parameters (Figure 5c-d). We further demonstrated that a stronger HSR activity – as predicted by the model simulations of the GFP slope – among isogenic nematode translates to a longer lifespan (Figure 6), validating the functional consequences of the two protein turnover parameters. Thus, our results point to the quality of maintenance of protein turnover (interconnection of translation and degradation rates) or more general to protein homeostasis (proteostasis), as major source for the heterogeneity in the length of the *C. elegans* adult lifespan.

The results of the mathematical modelling and parameter estimation of HSR dynamics in our study further suggest that the degree of age-related proteostatic decline varies among individual *C. elegans* and across compartments within an animal. The discrepancy in the proteostatic decline has the effect of increasing animal-to-animal heterogeneity in the protein turnover and HSR dynamics across the whole-body as well as within individual compartments of the nematodes. By contrast, the protein turnover parameters and HSR capacity along the body of a single animal become more alike. Taken together, the results point to a higher inter-animal heterogeneity but lower intra-animal variability of proteostasis and stress response capacity with aging.

An accumulation body of evidence suggests that in somatic tissues of *C. elegans* proteostasis collapses after reaching adulthood [52]. This may be due to allocating resources to the growth of embryos as postulated by the disposable soma theory of aging. For instance, in early adulthood, the *C. elegans* intestine starts producing yolk proteins in high amounts that are secreted and picked up by the developing embryos [61]. Hence, intestinal cells can re-allocate their resources by maintaining the intestinal proteome, which includes the expression of molecular chaperones to production of yolk proteins. Indeed, we observed the decline of HSR in the intestine as the most significant in magnitude among the compartments (Supplementary Table 4 and 5), as well as the most stochastic (*i.e.*, the highest increase in animal-to-animal variability), We speculate that the decision from investment in protein maintenance to production of yolk underlies some stochastic components of fluctuating expression levels of key regulators, analogues to the observed stochasticity of the decision making of bacteria phage lambda to enter the lysogenic or lytic cycle [60]. Interestingly, inhibiting genes in the yolk production and conversely overexpressing chaperone HSP-16 is both sufficient to increase lifespan of *C. elegans* [62, 63], suggesting that interfering with the expression these genes might tip the scale towards somatic protein maintenance, resulting in systemic effects on organismal longevity.

## Methods

### a. *C. elegans* strains and culture

*C. elegans* populations were cultured and maintained at 20° C on nematode growth media (NGM) plates and fed with *E. coli* OP50 using standard protocols [64] before and between microfluidic experiments. Heat-shock was delivered to the animals by placing the NGM plates with *C. elegans* inside an incubator at 37° C for one hour. Age-synchronized day-1 and day-2 adults, via bleaching, were used in experiments. For population and individual assays in microfluidic devices, animals were suspended in M9 solution before each experiment, repeatedly washed to ensure the complete removal of progeny and debris. The transgenic *C. elegans* strain TJ375 [P*hsp-16.2*∷GFP(gpIs1)] [20] was provided by the *Caenorhabditis* Genetics Center at the University of Minnesota. TJ375 features the 400-bp *hsp-16.2* promoter coupled to the gene encoding green fluorescent protein (GFP) but not encoding the protein HSP-16.2. Accordingly, this strain allows for the quantitative assessment of the amount of native HSP-16.2 in *C. elegans* following heat-shock when measuring its fluorescence intensity. We crossed and used *C. elegans* strains LSD2088 (*jmjd-3.1(gk384)*, P*hsp-16.2*∷GFP(gpIs1)) and LSD2089 (*ncl-1(e1942)*, P*hsp-16.2*∷GFP(gpIs1)). See primers in Supplementary Table 6.

### b. Microfluidic device fabrication

Microfluidic devices were fabricated using standard soft lithographic techniques and consisted of one structured polydimethylsiloxane layer bonded to a planar glass slide. Briefly, microfluidic circuits were designed using AutoCAD 2014 (Autodesk, San Rafael, USA) and printed onto a high-resolution film photomask (Micro Lithography Services Ltd, Chelmsford, UK). Master structures were fabricated on SU-8 (Microchem Corporation, Westborough, USA) coated silicon wafers using conventional photolithographic methods [65], with feature heights being 95 μm for the “high-throughput” devices (for adult *C. elegans*) and 65 μm for the “long-term immobilization” devices (for adult *C. elegans*). Microfluidic devices were subsequently manufactured using standard soft-lithographic techniques [66]. Specifically, a 20:1 wt/wt mixture of polydimethylsiloxane base and Sylgard 184 curing agent (Dow Corning, Midland, USA) [67] was poured over the SU8 mould structure and cured in oven at 70°C for 8 hours. Such a composition ensured that the PDMS was flexible enough for efficient PDMS-valve actuation. The cured PDMS layer was then peeled off, cut to shape and inlet/outlet were formed using a hole puncher. Finally, the structured PDMS layer was bonded to a planar glass slide (Menzel Gläser, Braunschweig, Germany) in an oxygen plasma, and the bonding was reinforced by a post-bake at 70° C for 3 hours.

### c. High-throughput *C. elegans* screening and sorting microfluidic devices

We developed two microfluidic devices for high-throughput screening and sorting of *C. elegans* based on their fluorescent protein reporter expression. The layouts of the devices are depicted in Figure 1e and Figure 2a, respectively. They consist of a central straight channel, where *C. elegans* (suspended in M9 buffer) flow in a continuous and non-overlapping manner. The nematodes enter the device through an inlet, which has wall-embedded PDMS blades and a central pillar to aid disaggregation of animals. Two additional buffer inlets enable appropriate spacing between single *C. elegans* flowing through the main channel. Whereas the screening device has only one outlet (Figure 1e), the sorting device has three outlets (Figure 2a), emanating from two consecutive bifurcations, subsequent to the main channel. In the sorting device (Figure 2a), during normal operation, one outlet is kept open to allow fluid and non-sorted *C. elegans* (*i.e.* animals expressing GFP levels lower than the minimum threshold value) to exit the device. Upon production of a sorting signal, the total flow is relocated through a specific outlet via actuation of lateral on-chip valves. These on-chip valves are formed by dead-end fluidic channels (Figure 2a, in pink) filled with deionised water and connected to high-pressure sources. Upon actuation, these valves acutely reduce the diameter (by approximately 95%) of the fluidic channel adjacent to the valve, increasing its hydraulic resistance and deflecting flow towards the remaining “open” channel. Finally, it should be noted that a reference channel is used for calibration, where fluorescence intensity originating from a standard dilution of fluorescein sodium salt (Sigma Aldrich, Buchs, Switzerland) is measured at a constant flow rate.

Efficient operation and performance of the high-throughput microfluidic devices were enabled by the functional integration of peripheral components of the system. These components are depicted in Supplementary Figure 2a,b. Continuous fluid flow was provided by precision syringe pumps (neMESYS, CETONI GmbH, Korbussen, Germany). Two dosing units delivered M9 buffer to the designated inlets of the device at volumetric flow rates of 50 and 70 μL/min respectively, while the suspension of *C. elegans* was controlled by other two fluid units (which delivered them at 80 μL/min). The optical system (Supplementary Figure 2a,b) provided a rapid detection of fluorescent protein reporters within animals. The optical system comprised a laser light source, an inverted confocal microscope and a photomultiplier tube (PMT) detector (Supplementary Figure 2b). For accurate *C. elegans* GFP protein quantification, a cylindrical lens was used to shape the laser beam into a light sheet orthogonal to the direction of *C. elegans* flow (an example of fluorescent signal read out is shown in Supplementary Figure 2e). A 488 nm laser beam (60 mW, Omicron Laser, Rodgau, Germany) was used as the excitation source and expanded using a telescope consisting of two plano-convex lenses and shaped into a light sheet using a cylindrical lens (LJ1558RM, *f* = 300 mm, Thorlabs, Newton NJ, USA). The light sheet was then directed into the back aperture of the microscope, using a dichroic mirror (AT DC 505, AHF, Tübingen, Germany) and subsequently focused into the microfluidic channel using a 20× objective (S Plan Fluor ELWD ADM, Nikon, Egg, Switzerland). Fluorescence emission was collected by the objective, passed through the same dichroic mirror and focused onto a 40 ± 3 μm pinhole (P40H, Thorlabs, Newton NJ, USA) through a tube lens (*f* = 200, Nikon, Egg, Switzerland). Subsequently, fluorescence was passed through an emission filter (525/45 nm, HC Bright Line Bandpass Filter, AHF, Tübingen, Germany) and onto a PMT (H10722520, Hamamatsu Photonics, Solothurn, Switzerland). A secondary detection system was used for visual inspection of device performance. Specifically, white light, bright field illumination filtered with a 610 nm emission filter (ET Longpass, AHF, Tübingen, Germany) was imaged using a high-speed camera (Phantom Miro, Vision Research, Baden-Baden, Germany). Using such a system, real-time images could be acquired in parallel without this (longer) wavelength light interfering with the fluorescence measurement. Collected light passed though the microscope beam splitter 80/20, with 20 % of the beam being directed to the high-speed camera, which was protected from the laser beam with a 532 nm longpass filter (LP Edge Basic, AHF, Tübingen, Germany) and 80 % being directed to the PMT. Sorting decisions were made by a FPGA (PCI NI-RIO, National Instruments, Austin, USA) (Supplementary Figure 2a) enabled system, and animal sorting was enabled by valve actuation, which promptly changes the fluidic resistance at given positions within the fluidic path, and directs the flow into a specific outlet. On-chip valves were connected to solenoid valves (MHA1, Festo, Esslingen, Germany), which were controlled by sorting impulses delivered by the FPGA. Valve-actuation times were defined by decision-making and the signal-output of the FPGA, the switch of configuration of the solenoid valve (approximately 4 ms) and the inflation time of the PDMS valve (between 8 and 15 ms). The total processing time for one *C. elegans* (from the fluorescence detection to the sorting in the proper outlet, enabled by actuation of valves) is 75 ms.

### d. Long-term *C. elegans* immobilization microfluidic device

The long-term immobilization microfluidic device consists of a main channel, whose size and shape allows efficient, yet gentle, immobilization of *C. elegans* (Figure 3a, in red) [30]. Two on-chip hydraulic valves positioned alongside the main channel allow control over nematode loading and immobilization (Figure 3a, in black). Hydraulic valves are formed by filling dead-end channels with water. Once pressurized, the wall separating the fluidic channel and valve channel deforms resulting in partially blockage of the fluidic channel. In devices designed for immobilization of adult hermaphrodite *C. elegans*, egg laying is facilitated through a set of pillars roughly positioned alongside the animal’s vulva. A highly concentrated bacteria suspension is supplied to an immobilized *C. elegans* as food source (Figure 3a, in yellow). *C. elegans* immobilized in such a device remain viable for more than 100 hours on average and exhibit normal physiological functions, with both feeding and egg laying occurring at rates comparable to free crawling *C. elegans* on NGM plates (Figure 3b). Furthermore, *C. elegans* are immobilized such that high-resolution fluorescence images can be acquired. These capabilities allowed quantitative long-term monitoring of the GFP expression after heat-shock in a non-invasive manner for individual animals.

All images were acquired on an inverted microscope (Nikon Ti-*S*, Nikon Inc.) equipped with a sCMOS camera (Prime 95B, Photometrics Inc.), a fluorescence LED (LedHUB, Omicron-laserage Laserprodukte GmbH) and a piezo objective drive (Nano-F100S, Mad City Labs Inc.), controlled by a custom build Matlab script (Matlab 2016b. Mathworks Inc.). Images of the immobilized *C. elegans* were acquired at 1-hour intervals, at each time-point a fluorescent Z-stack was acquired (26 slices with 2 μm spacing) using an exposure time of 10 ms and 50 % excitation intensity. All images were acquired using a 10× objective (CFI Plan Fluor 10×, Nikon Inc.). We quantified the mean fluorescence intensity from each *C. elegans* at each time point-measuring a defined region of interest which comprises the whole animal-with the open source software package ImageJ (NIH, Bethesda, USA).

### e. Long-term *C. elegans* embryo microfluidic device

The long-term *C. elegans* embryo microfluidic device consisted of a simple straight channel within which embryos were confined for long-term imaging (Figure 8a). Such a channel was designed with an initial width of approximately 50 μm, so that single embryos could flow into the device and would remain on-chip once loading was completed. Following the loading region, the channel expanded to 200 μm width, here the channel width was chosen such that the channel would fit into the microscope field of view at 40-60X magnification. An array of small pillars was placed into the wide channel, trapping embryos within the channel. Channel height was chosen at 30 μm, such that most of the embryos would be oriented for optimal imaging (Figure 8b).

Embryos were harvested from adult hermaphrodite animals, at different ages (Adult day-1, day-2 and day-4). Following heat-shock at 37°C for 1-hour, adult animals were synchronized via bleaching (200 μl of 1 M NaOH and 400 μl of 10% NaClO), the embryos collected and washed twice with M9 buffer. Embryos were passed through a 40μm cell strainer to remove any debris, loaded into a short piece of tubing and connected to the device. Prior to loading, the microfluidic device was filled with M9 buffer and gently pressurized to remove all air from the device. Embryos were finally loaded into the device by gently applying pressure to the syringe attached to the embryo inlet, and passively trapped on the pillar array placed into the wide imaging channel. Thirty Z-stacks of the embryos throughout the heat shock response were recorded on an epi-fluorescence microscope with a spacing of 1 μm using a 40× oil immersion objective. Stacks were acquired at 10 minute intervals, for a total duration of 12 hours. We quantified the mean fluorescence intensity from each *C. elegans*’s embryo at each time point-measuring specific regions of interest which comprised a single embryo-with the open source software package ImageJ (NIH, Bethesda, USA) (Figure 8c).

### f. Western blot quantification of GFP and HSP-16.2

Western blot quantification of native GFP and HSP-16.2 was carried out in strain TJ375 in order to study their relative expression at different time points after heat-shock and validate our measurements. For this, samples of *C. elegans* strain TJ375 were harvested at different time points (4, 8, 12, 18, 24, 48 h) after heat-shock, as well as TJ375 samples without heat-shock. N2 control samples were harvested 8 h after heat-shock and without heat-shock (Figure 1c,d and Supplementary Figure 1). Around 2 000 *C. elegans* per sample were placed in M9, frozen immediately in liquid nitrogen and maintained at −80° C. Protein extraction from *C. elegans* samples was done by adding 150 μl lysis buffer (RIPA buffer (ThermoFisher #89900, Reinach, Switzerland), 20 mM sodium fluoride (Sigma #67414, Buchs, Switzerland), 2 mM sodium orthovanadate (Sigma #450243, Buchs, Switzerland), and protease inhibitor (Roche #04693116001, Basel, Switzerland)) to each *C. elegans* pellet and subsequently sonicating them repetitively until tissues were dissolved. Aliquots were collected and processed. For equal loading the protein concentration of the supernatant was determined with BioRad DC protein assay kit II (BioRad#5000116, Cressier, Switzerland) and standard curve with Albumin (ThermoFisher #23210, Reinach, Switzerland). After 5 minutes of boiling at 95° C, 10 μg of protein in 20 μl of loading buffer (42.5 mM Tris-HCl pH 6.8, 1.7% SDS, 8.5% glycerol, 15% 2-mercaptoethanol, and 1% bromophenol blue, final concentrations) were loaded per sample in each well of a NuPAGE 10 % Bis-Tris protein gel (ThermoFisher #NP0301BOX, Reinach, Switzerland). Gel electrophoresis was done for 1.5 h at 130 V using a Mini Gel Tank filled with MOPS 1× buffer. Right after, the gel was electro-blotted onto a nitrocellulose membrane (Sigma #GE10600002, Buchs, Switzerland) using a Mini Trans-Blot Cell filled with transfer buffer (25 mM Tris-Base, 0.2 M glycine and 20#x0025; methanol, final concentrations) and applying 100 V for 1 h. Subsequently, the membrane was blocked for 1 h using 5% milk powder in TBS-t buffer (50 mM Tris-Base, 0.15 M NaCl and 0.1% Tween-20, pH 7.2-7.6) at room temperature. Right after, the membrane was cut (according to bands of molecular weight indicated by the protein ladder) in 3 pieces for identification of HSP-16.2, GFP and Tubulin respectively. The membrane for identification of HSP-16.2 was incubated in 5% milk powder in TBS-t buffer with a polyclonal anti-HSP-16.2 primary antibody (1:8000) courtesy of Prof. Gordon Lithgow. The membrane for identification of GFP was incubated in 5% milk powder in PBS-t buffer (0.13 M NaCl, 2.7 mM KCl, 10 mM Na_2_HPO_4_, 1.8 mM KH_2_PO_4_ and 0.2% Tween-20, pH 7.2-7.6) with anti-GFP primary antibody (1:32000, Roche #11814460001, Basel, Switzerland). Lastly, the membrane for identification of Tubulin, as control protein, was incubated in 5% milk powder in TBS-t buffer with anti-tubulin primary antibody (1:500, Sigma #T9026, Buchs, Switzerland)). All membranes with primary antibodies were incubated overnight at 4° C. Then, after through TBS washes, blots were probed with secondary antibodies: Tubulin and GFP with anti-mouse IgG antibody (1:2000) and HSP-16.2 with anti-rabbit IgG antibody (1:1500) in TBS-t suspensions with 5% milk powder for 1 h and 1.5 h respectively. Finally, the protein bands were visualized upon few minutes of incubation of the membranes with Bio-Rad Clarity Western ECL Substrate (Bio-Rad#1705061, Cressier, Switzerland). Blots were imaged using ChemiDoc imaging system for chemiluminescence assays with 0.1 s exposure. Density of the bands was quantified using software package ImageJ (NIH, Bethesda, USA), and normalized to Tubulin values (Figure 1c,d and Supplementary Figure 1).

### g. HSR mathematical modelling

#### Model description

The mathematical model of HSR consists of 12 species (Table 1), 15 reactions (Figure 4a & Table 2) – five of which are reversible – with 24 parameters. The full model equations are provided in Supplementary Data File 2. The model simulates the cellular response to an increase in intracellular misfolded protein caused by a heat-shock. Prior to the simulation of heat shock response, we performed a pre-equilibration step by running the model until steady state (*i.e.* until the concentrations stabilize). Heat shock stress was simulated by increasing the misfolding rate of proteins (*Prot*) from *α* to *β* (*β* > *α*) for a period of 1 hour, emulating the transfer of *C. elegans* from a 20° C to a 37° C environment. In the model, misfolded proteins (MFP) are captured by molecular chaperons, specifically the heat shock protein (HSP), for protein refolding. As a consequence of the heat-shock, Heat Shock Factor 1 (HSF-1) molecules that would otherwise form a complex with HSP (HSP:HSF-1) become free. Free HSF-1 molecules subsequently undergo reversible dimerization (HSF-1_2_) and trimerization (HSF-1_3_). HSF-1 trimers are able to translocate to the nucleus, bind to the heat shock element (HSE), and promote the expression of the HSP. The newly produced HSP (in excess of the level needed to refold MFP) will bind to free HSF-1 molecules and thereby repress its own expression. For the TJ375 strain, GFP is encoded downstream of a promoter containing an HSE. Accordingly, we modified the original HSR model to include a balance equation for the GFP, in which the expression rate is the same as that of HSP. As a reporter, the GFP does not interact with any other species in the model, and thus does not influence the HSR dynamics. It should be noted that the model does not include a balance equation for the protein and HSF-1, and thus the total amount of protein (*Prot* + MFP) and HSF-1 does not vary with time. However, we allowed the amount of protein (*Prot*) and HSF-1 to differ among individual *C. elegans* (see below).

#### Parameter estimation for the average HSR dynamics of adult C. elegans

In the first stage of parameter estimation, we obtained the model parameter estimates by fitting the simulated GFP concentration to the average fluorescence values amongst all measured *C. elegans*. When comparing model prediction of GFP expression and the average fluorescence data, we scaled the value of the simulated GFP concentration such that its peak equals to the maximum value of the average fluorescence intensity. The parameter estimation was formulated as a minimization of the objective function:

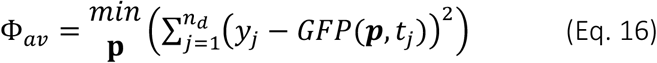

where ***p*** is the model parameter vector, *n*_*d*_ is the number of measurement data (*n*_*d*_ = 48), *y*_*j*_ is the average GFP fluorescence intensity among the *C. elegans* at time *j*, and *GFP*(*p*,*t*_*j*_) is the simulated GFP concentration at time *j* for the parameter vector ***p*** The lower and upper bounds of the parameter ***p***, denoted by *lb*_*k*_ and *ub*_*k*_ (*i.e. lb*_*k*_ ≤ *p*_*k*_ ≤ *ub*_*k*_) for the *k*-th parameter, are set to 2 orders of magnitude below and above those of the original model parameter values reported in Scheff *et al.* [43]. The optimization was solved using a hybrid scatter search algorithm called MEIGO in MATLAB [68]. The resulting parameter estimates are provided in Table 3.

#### Sensitivity analysis

We performed a local sensitivity analysis of the model using the parameter estimates in Table 3, by perturbing each parameter one-at-a-time and simulating the resulting GFP expression curve. We then calculated the sensitivity coefficients of the HSR dynamics for the following: the maximum value of GFP and the time to reach the GFP maximum value. In this manner, we identified the parameters that exert a strong influence on the slope of the GFP expression curve. The sensitivity coefficients were computed using a finite difference approximation, as follows:

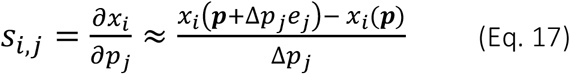

where *s*_*i*,*j*_ is the sensitivity coefficient of *x*_*i*_ with respect to perturbation to the *j*-th parameter, *x*_*i*_ is one of the two metrics of HSR dynamics above,Δ*p*_*j*_ denotes the perturbation to the *j*-th model parameter – chosen to be 10 % of the nominal parameter values - and *e*_*j*_ is the unit basis vector in the *j*-th direction. The result of such a sensitivity analysis is presented in Supplementary Figure 6. The sensitivity analysis suggests that the parameters related to the HSP (GFP) transcription and translation, specifically *HSE*_*0*_, *k*_*4*_, *k*_*5*_, *k*_*6*_, and *k*_*11*_, the amount of proteins *HSF*_*0*_ and *Prot*_*0*_, and not surprisingly the protein misfolding rates *α* and *β*, have the strongest influence on the GFP dynamics before its peak.

#### HSR dynamics in single C. elegans

We further adapted the HSR model of the average *C. elegans* above to describe the HSR time course from single *C. elegans*. We kept the ODE model formulation the same as described above, and further assumed that individual *C. elegans* differ from the average *C. elegans* only for a few model parameters. More specifically, for the single *C. elegans* parameter fitting, we only considered parameters related to biologically regulated process such as the protein turnover and set the other physicochemical parameters such as protein misfolding rates and HSF-1 dimerization/trimerization to be constant across different animals (see Table 3). The results from the model sensitivity analysis above point to the regulation of protein amount as the most important process that modulates the HSR dynamics. With these considerations, we narrowed down the possible model parameters that vary among animals to *k*_*4*_, *k*_*5*_, *k*_*6*_, and *k*_*11*_ and the initial amount of proteins *HSF*_*0*_ and *Prot*_*0*_. Note that since the model does not include new synthesis and degradation of any forms of *HSF* and *Prot* by assuming that they exist at pseudo steady state level, the initial amount of *HSF*_*0*_ and *Prot*_*0*_ thus reflects the total amount at any time.

Among the six parameters, four are related to protein turnover, specifically the protein translation and degradation rate constants for HSP (GFP) and the total concentration of proteins *HSF*_*0*_ and *Prot*_*0*_. Also, there exists a time-scale separation between the mRNA and the protein HSP (GFP) with the mRNA having much faster dynamics. Importantly, comparing TJ375 with LSD2088, a strain with mutant histone demethylase *jmjd-3.1* causing alterations in the chromatin accessibility of HSF-1 regulation of HSP, the heterogeneity of HSR remained relatively unchanged (see Supplementary Figure 4). Thus, in the modelling of HSR dynamics in single animals, we vary the protein turnover rates for each *C. elegans* by scaling the translation and degradation rates of HSP as well as the amount of proteins *HSF*_*0*_ and *Prot*_*0*_. The total amount of proteins *HSF*_*0*_ and *Prot*_*0*_ in individual *C. elegans* scales with the parameters *k*_*6*_ and *k*_*11*_ as follows: (see derivations in Supplementary Data File 3)

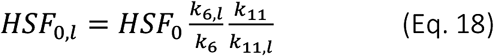

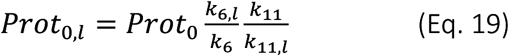

where the subscript *l* refers to the parameters from the specific *C. elegans l* and the non-subscripted parameters are taken from the average *C. elegans* estimates in Table 3.

We performed the single *C. elegans* parameter fitting in the same manner as for the average *C. elegans* except for the fluorescence data, which now came from individual *C. elegans*:

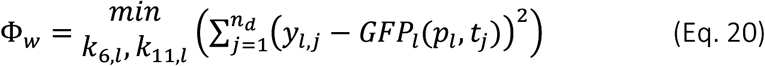

where *n*_*d*_ is the number of measurement data (*n*_*d*_ = 48), *y*_*l*,*j*_ is the measured GFP intensity of *C. elegans l* at time *j* and *GFP*_*l*_ is the simulated GFP concentration at time *j* for that specific *C. elegans l*. The upper and lower bound of the parameters were allowed to vary by 1 order of magnitude from the values in Table 3. This optimization was performed using the MATLAB function *fmincon* with multiple starts (*n* = 120). The results obtained from the parameter estimation of individual *C. elegans* fluorescence data are shown in Figure 4d.

#### HSR dynamics in embryo C. elegans

Finally, we expanded our mathematical model for describing HSR in embryo *C. elegans*. Keeping the ODE model formulation previously described, we used the time-course HSR experimental data obtained from embryo *C. elegans*. We first obtained the model parameter estimates by fitting the simulated GFP concentration to the average fluorescence values amongst all measured *C. elegans* eggs, shown in Figure 8c. The optimization was as well solved using MEIGO in MATLAB [68], by iterating 1000 times the calculation, starting from the values shown in Table 3, which describe HSR in adult *C. elegans*. The result is shown in Table 4.

Subsequently, we found the individual translation and degradation constant rates (*k*_*6*_ and *k*_*11*_) which would describe the individual time course HSR of individual embryos. For this, we reproduced the procedure described in the previous section (*HSR dynamics in single C. elegans*). The resulting parameter estimates for individual embryo *C. elegans* are shown in Figure 8d and Supplementary Table 7.

### h. Lifespan machine

The automated lifespan analysis was performed using the lifespan machine platform described by [56]. The experiment was prepared identically to the manual lifespan setup with the exception of using tight-fitting petri dishes (BD Falcon Petri Dishes, 50×9mm) and drying the plates without their lids for 30 minutes prior to starting the experiment. The automated analysis was performed in air-cooled Epson V800 scanners using a scanning frequency of one scan per 30 minutes. The temperature was monitored using temperature probes (Thermoworks, Utah, US) on the scanner flatbed and kept constant at 20° C.

### i. Manually performed lifespan (by hand)

Manual lifespan scoring (by hand) was performed as described in [69]. In brief, about 2000 TJ375 animals were heat-shocked and sorted as described above and then about 100 L4 *C. elegans* per sorting category were picked on NGM plates containing 50 μM 5-Fluoro-2’deoxyuridine (FUDR), 15 μg/ml nystatin and 100 μg/ml carbenicillin seeded with heat-killed OP50 bacteria [70] and scored at 20°C. Animals were classified as dead if they failed to respond to prodding. Exploded, bagged, burrowed, or animals that left the agar were censored. Population survival was analyzed with the statistical software R [71] using the survival [72] and survminer [73] packages and with the L4 stage defined as timepoint zero. The estimates of survival functions were calculated using the product-limit (Kaplan-Meier) method and the log-rank (Mantel-Cox) method was used to test the null hypothesis.

## Acknowledgements

We thank Gordon Lithgow for the generous gift of the HSP-16.2 antibody. Some strains were provided by the CGC, which is funded by NIH Office of Research Infrastructure Programs (P40 OD010440). We thank Jan Gruber for his valuable contributions in the writing of the original SNSF grant and discussions. We also acknowledge Richard Venz for his valuable help in the western blot assays and creation of the new *C. elegans* strains, as well as Oliver Dressler and Ankit Jain for their help in the Labview coding of the sorting algorithm. Finally, we acknowledge Benjamin Towbin for his critical reading of the manuscript.

This research was funded by the Swiss National Science Foundation PP00P3_163898 to CYE and CS, and 2-77716-13 C-Elegans for NVQ, SB and XCS.

## Authors contributions

N. V.-Q., X. C. i S., C. Y. E., R. G. and A. dM. conceived, designed and interpreted the work presented in this work, N. V.-Q. and S. B. developed, conducted and analysed the experiments using microfluidic platforms, X.C. i S. and S.S. contributed to the construction of experimental microfluidic setup, C. S. conducted the lifespan experiments, J. A. and P. R. worked on the mathematical modelling and N. V.-Q., C. Y. E., R. G. and A. dM. drafted and critically reviewed the manuscript

