## SupplementaryInformation for "Heterogeneity in heat shock response dynamics caused by translation fidelity decline and proteostasis collapse"

### Supplementary Information

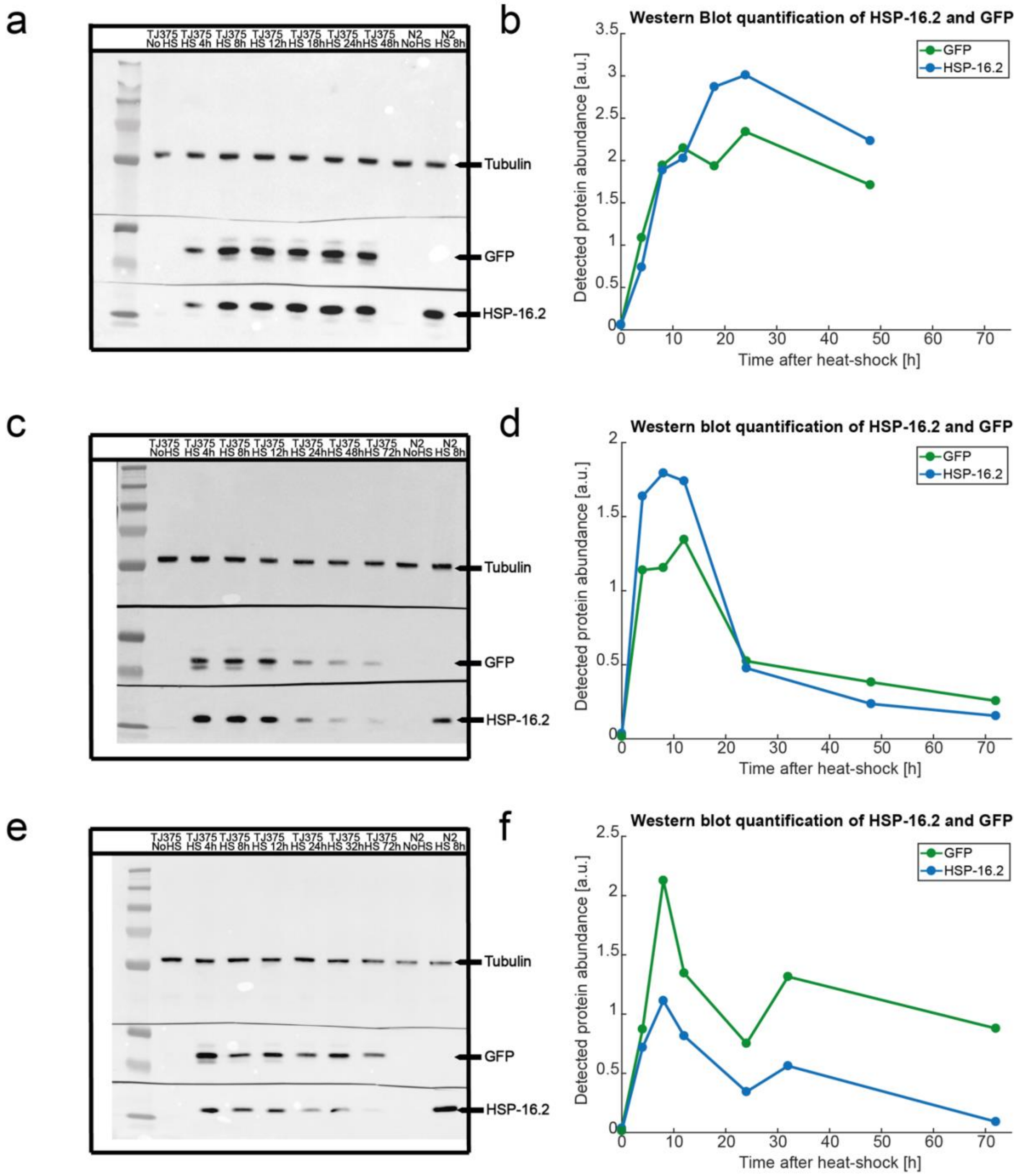


Supplementary Figure 1. Western Blots for protein detection and quantification of GFP and HSP-16.2. (a,c,e) Images of 3 biological repeat blots of 9 *C. elegans* samples (7 TJ375 samples at different times after heat-shock and 2 N2 with and without heat-shock). In all assays we use the housekeeping protein Tubulin as reference for relative protein quantification. (b,d,f) Corresponding quantifications of GFP and HSP-16.2 for samples at different times after heat-shock. A direct relation between GFP and HSP-16.2 abundance in these times is observed.

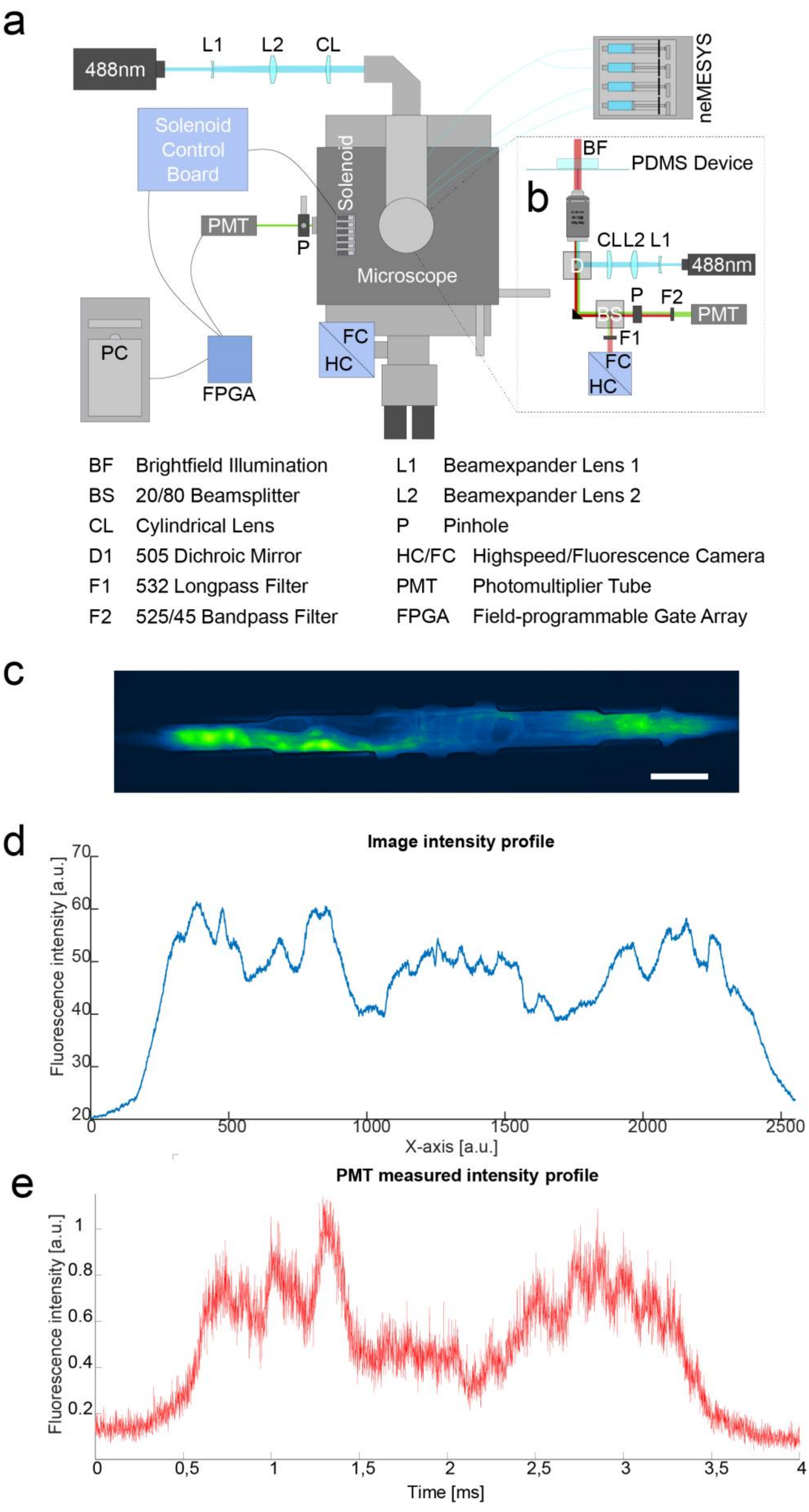

a
488nm
L1
L2
CL
neMESYS
Solenoid Control Board
Solenoid
PMT
P
Microscope
PC
FPGA
FC
HC
BF
PDMS Device
b
CL L2 L1
488nm
D
P
F2
BS
PMT
F1
FC
HC
BF Brightfield Illumination
BS 20/80 Beamsplitter
CL Cylindrical Lens
D1 505 Dichroic Mirror
F1 532 Longpass Filter
F2 525/45 Bandpass Filter
L1 Beamexpander Lens 1
L2 Beamexpander Lens 2
P Pinhole
HC/FC Highspeed/Fluorescence Camera
PMT Photomultiplier Tube
FPGA Field-programmable Gate Array
c
d
Image intensity profile
Fluorescence intensity [a.u.]
X-axis [a.u.]
0
500
1000
1500
2000
2500
20
30
40
50
60
70
e
PMT measured intensity profile
Fluorescence intensity [a.u.]
Time [ms]
0
0,5
1
1,5
2
2,5
3
3,5
4
0.2
0.4
0.6
0.8
1

**Supplementary Figure 2. High-throughput microfluidic system for *C. elegans* fluorescent proteins screening and sorting.** (a-b) Schematic representation of the complete experimental setup for sorting of *C. elegans* populations based on analysis of fluorescent protein reporters. (a) The microfluidic device is placed onto the stage of an inverted microscope. Four dispensing NemeSYS syringe pumps control the flow rate delivered to the microfluidic device. A 488 nm laser beam is expanded and shaped with a cylindrical lens for illuminating the sample in the form of a light sheet. The emitted light from the sample is collected and directed to a PMT and shown in detail in panel (b). A high-speed camera is used for visual inspection of device operation. The actuation of on-chip valves is enabled by 2-way solenoid valves, whose control board is connected to a FPGA. The FPGA is connected to a PC where, through a custom LabVIEW software, the user can set the parameters for *C. elegans* sorting. (b) Optical setup for fluorescent proteins excitation and emission collection onto a PMT. (c-e) Comparison of the quantification of GFP expression in *C. elegans* with image analysis software (ImageJ) and with our optical system. (c) Fluorescence image of an immobilized animal heterogeneously expressing GFP. Scale bar represents 100 µm, image presented with ImageJ lookup table “green fire blue”. (d) Intensity profile of the fluorescent image, along its x-axis, presented in (c), measured with ImageJ. (e) Fluorescence signal readout obtained with the optical system (b) of the animal shown in (c).

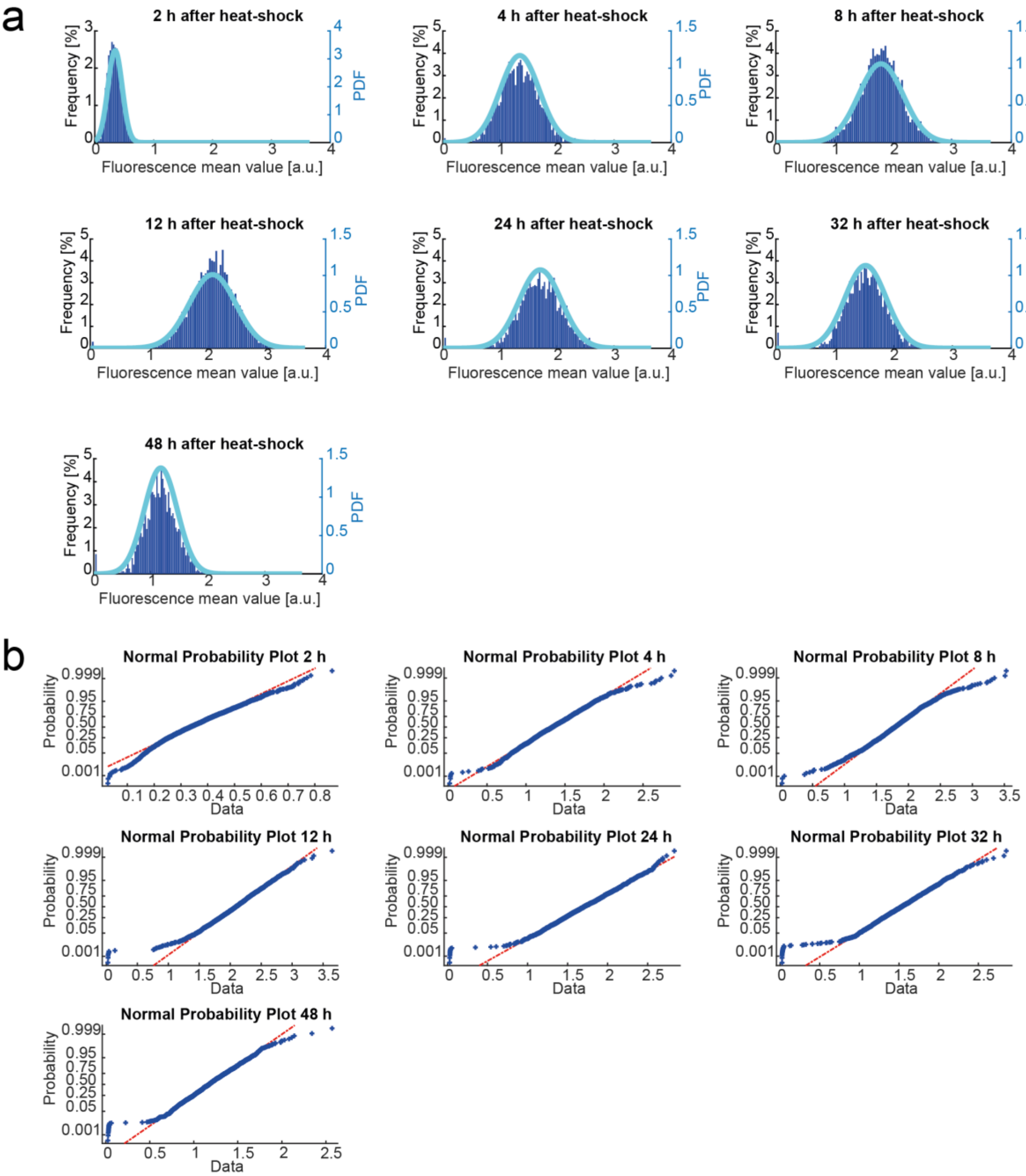


c

| | TJ375 adult day-2 | | |
|---|---|---|---|
| **Time** | **Mean** | **Std. Dev.** | **CV** |
| 2 h | 0,344 | 0,121 | 35,2 % |
| 4 h | 1,348 | 0,341 | 25,3 % |
| 8 h | 1,786 | 0,377 | 21,1 % |
| 12 h | 2,078 | 0,395 | 19,0 % |
| 24 h | 1,707 | 0,371 | 21,7 % |
| 32 h | 1,525 | 0,351 | 23,0 % |
| 48 h | 1,173 | 0,289 | 24,6 % |

Supplementary Figure 3. GFP mean values expression distribution in a *C. elegans* population at different time points after heat-shock. (a) Distribution of GFP mean expressed values in a *C. elegans* population at different time points (2, 4, 8, 12, 24, 32 and 48 hours after heat-shock). Overlaid on each histogram we show the normal probability distribution function (PDF) at each time point. (b) Normal probability plots of the *C. elegans* population shown in (a). Theoretical distributions are shown with the red lines and the blue crosses are the actual data points. (c) Mean, standard deviation and coefficient of variation values of the GFP expression on *C. elegans* populations shown in (a). All values presented are in a.u.

a

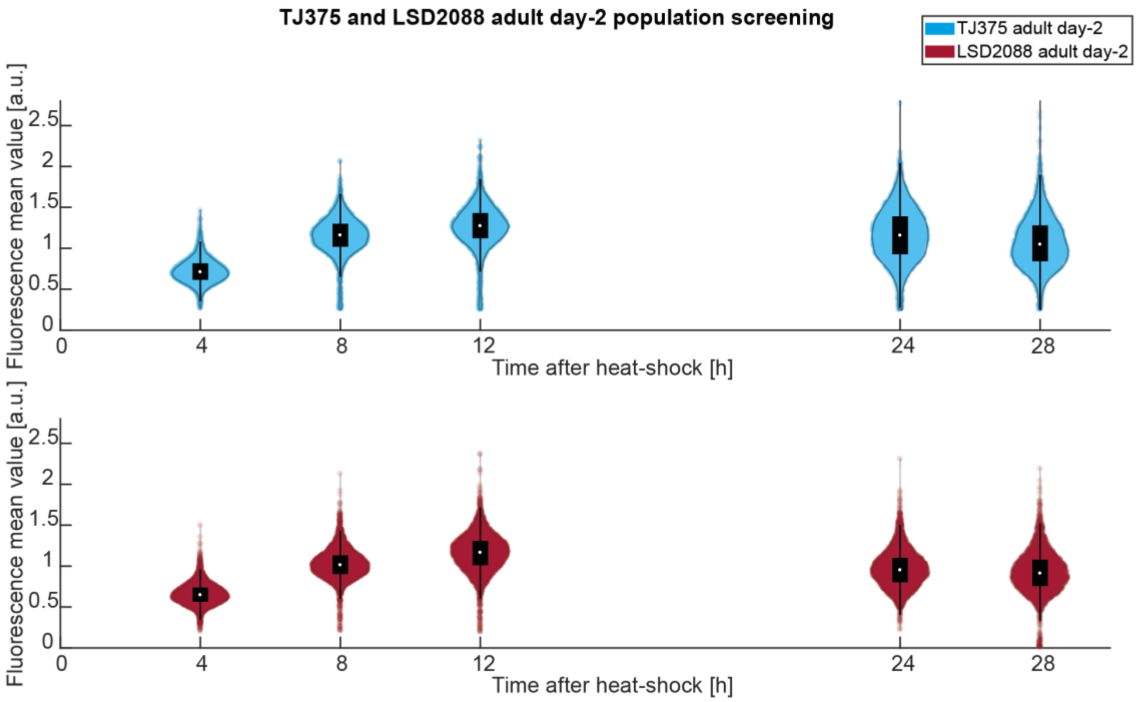


b

| | TJ375 adult day-2 | | | LSD2088 adult day-2 | | |
|---|---|---|---|---|---|---|
| Time | Mean | Std. Dev. | CV | Mean | Std. Dev. | CV |
| 4 h | 0,72 | 0,15 | 20,7 % | 0,66 | 0,13 | 19,5 % |
| 8 h | 1,15 | 0,23 | 19,6 % | 1,02 | 0,18 | 17,2 % |
| 12 h | 1,26 | 0,26 | 20,6 % | 1,16 | 0,23 | 19,5 % |
| 24 h | 1,16 | 0,33 | 28,0 % | 0,96 | 0,22 | 22,8 % |
| 28 h | 1,08 | 0,32 | 29,5 % | 0,93 | 0,25 | 26,7 % |

**Supplementary Figure 4. GFP longitudinal mean expression distributions in *C. elegans* populations at different time points after heat-shock (TJ375 and LSD2088 strains).** (a) Violin plots of populations TJ375 (P*hsp-16.2*::GFP) and LSD2088 (*jmjd-3.1*, P*hsp-16.2*::GFP) at different time points (4, 8, 12, 24 and 28 hours) after heat-shock, measured with the high-throughput microfluidic system presented in Figure 1e. (b) Mean, standard deviation and coefficient of variation (CV) of the expressed GFP in the populations shown in (a).

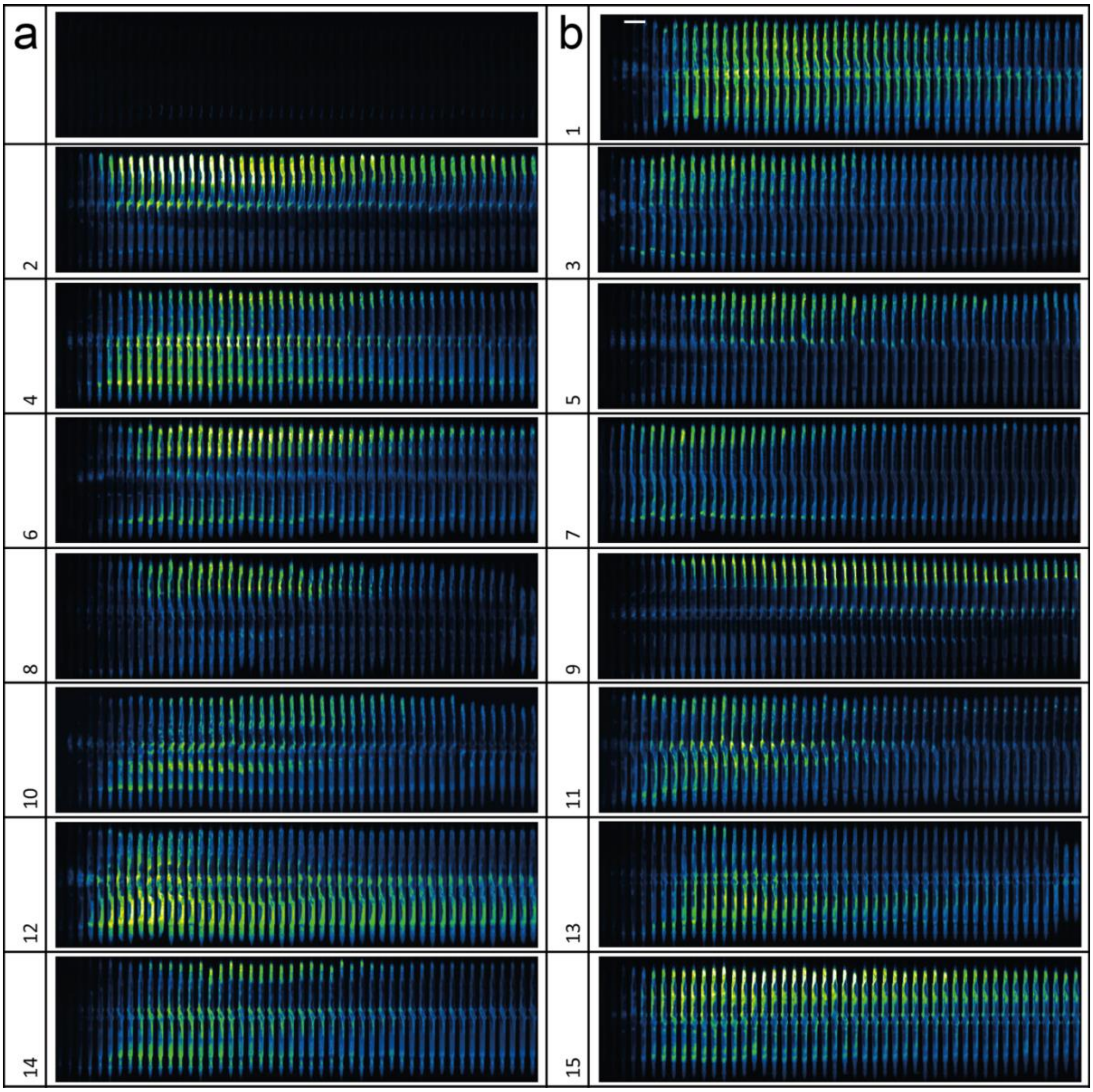


**Supplementary Figure 5. Image time-series of individual TJ375 *C. elegans* over 48 hours after heat-shock.** (a-b) Fluorescent images from immobilized TJ375 *C. elegans* in a microfluidic device. In each panel 48 images are shown, taken each progressively with 1-hour difference. In all panels the image on the left-hand side is taken after 1 hour and the image on the right-hand side is taken after 48 hours. In all panels the animals are shown with the head down. (a) *C. elegans* without experiencing heat-shock. (b) 15 adult day-2 *C. elegans* after heat-shock. To be noted, the heterogeneity of the HSR is seen in tissues expressing GFP and the temporal variation of this expression among them. Bar scale represents 200 µm. All images are shown with the ImageJ lookup table “green fire blue”.

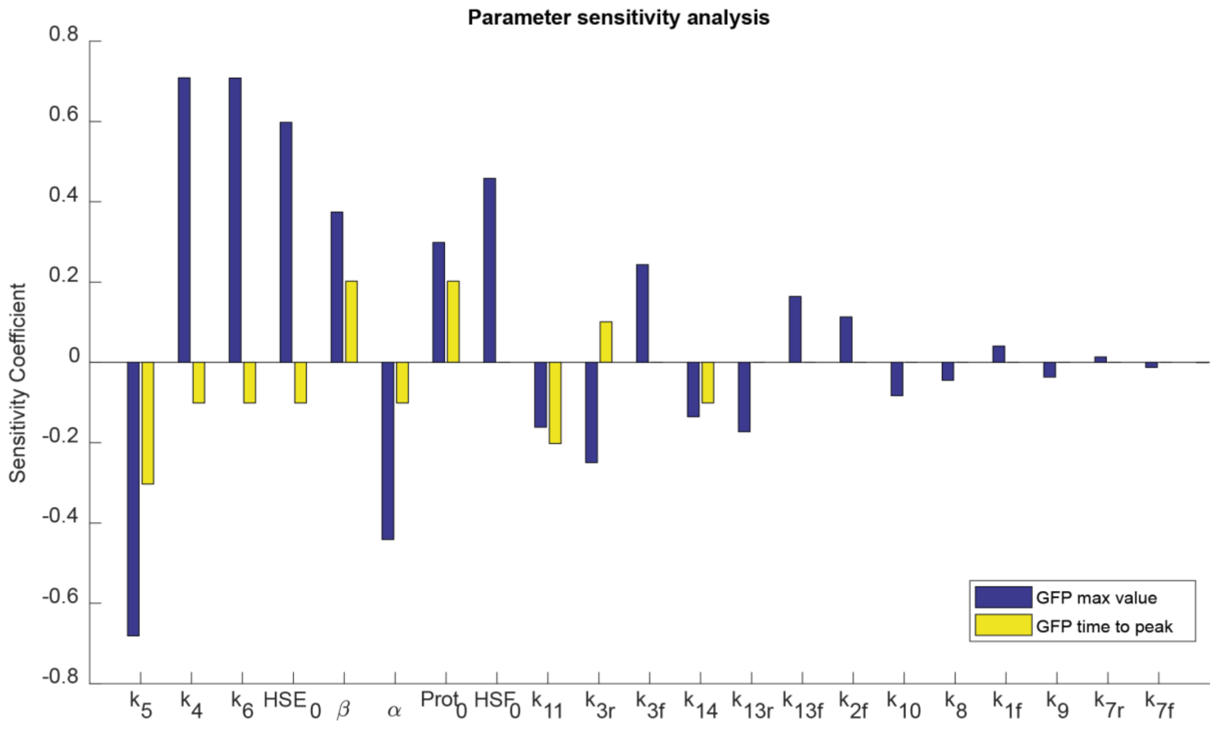


Supplementary Figure 6. Sensitivity values for the mathematical model parameters. Ordered from highest to lowest, we displayed the parameters with their sensitivity coefficients, with respect to the two metrics that describe the initial slope of the GFP expression curve, that is the fluorescence maximum value (GFP peak) and the time to achieve this fluorescence maximum value (GFP rise time). See Table 1 for species full names and Methods for details.

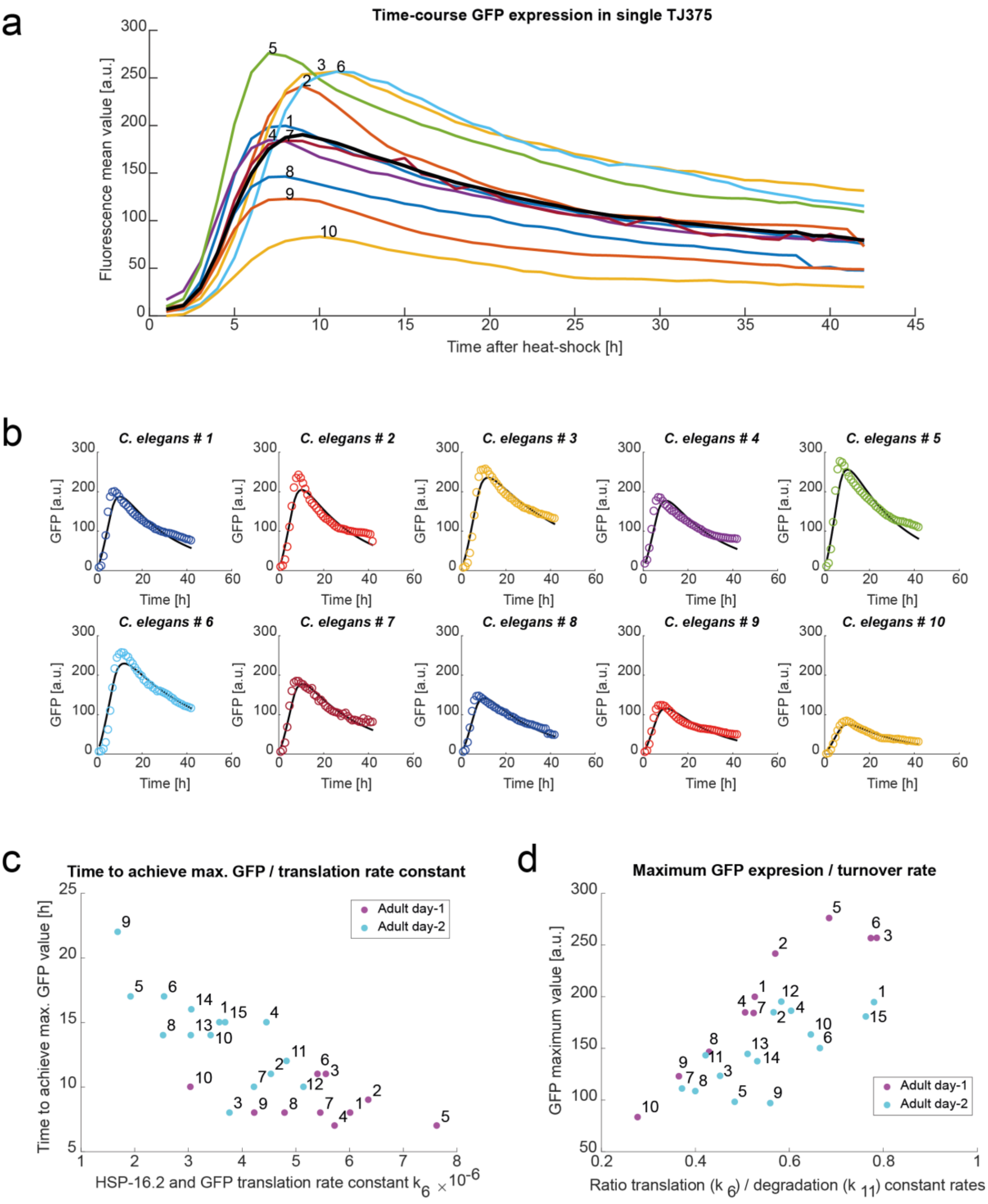


**Supplementary Figure 7. Adult day-1 GFP expression curves and parameter analysis of their influence in expression curves.** (a) Measured GFP mean expression over time ($t$ = 42 h) in individual adult day-1 TJ375 *C. elegans*. Each curve represents a single *C.elegans* ($n$ = 10) and they are identified with a number next to the curve. (b) Individual fitting of individual GFP expression for each analysed animal shown in (a), by varying only parameters $k_6$ and $k_{11}$ from the average response. Here the solid lines represent the modelled HSR while the circles represent the experimental data. (c) Correlation between the time to achieve the maximum GFP value for each *C. elegans* and its GFP translation rate constant ($k_6$). The time needed for individual response curve to achieve the maximum level of GFP expression in each animal is inversely proportional to the protein translation rate constant $k_6$. (d) Correlation between maximum GFP mean value for each *C. elegans* and its protein turnover. The maximum values of GFP concentration are positively correlated with the ratio between protein translation and degradation rate constants ($k_6/k_{11}$).

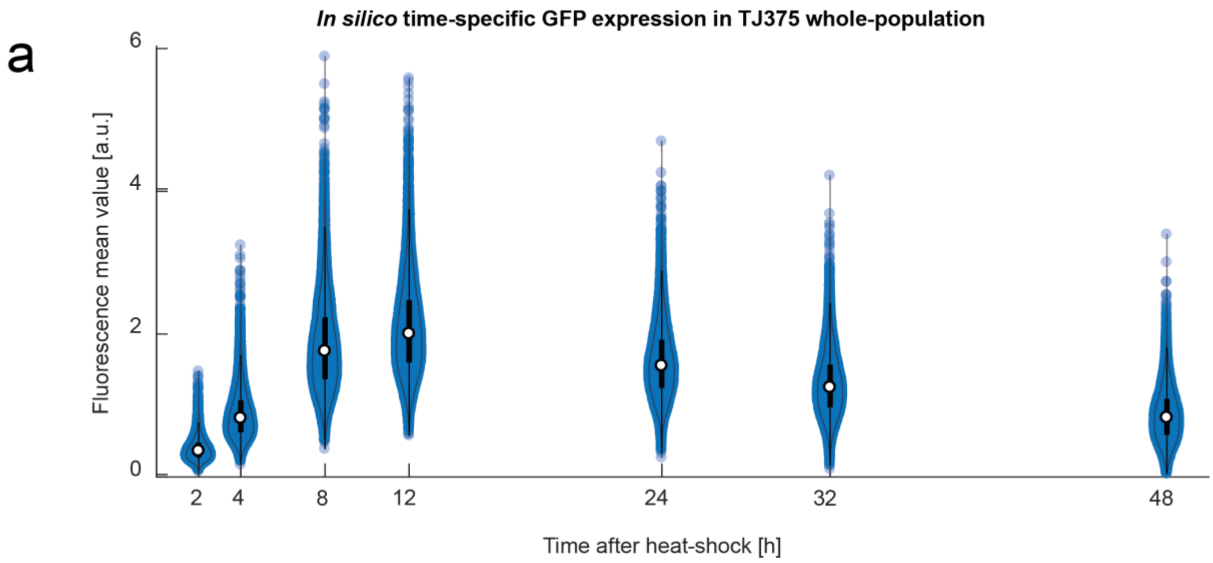


**Supplementary Figure 8. Comparison between simulated and measured GFP expression values of a TJ375 adult day-2 population after heat-shock**. (a-b) Violin plots of the fluorescence mean values of individual *C. elegans* TJ375 simulated (a) and measured (b) in a population at determined time points (2, 4, 8, 12, 24, 32 and 48 hours) after heat-shock. The thicker lines inside the violins represent 50 percent of the population.

a

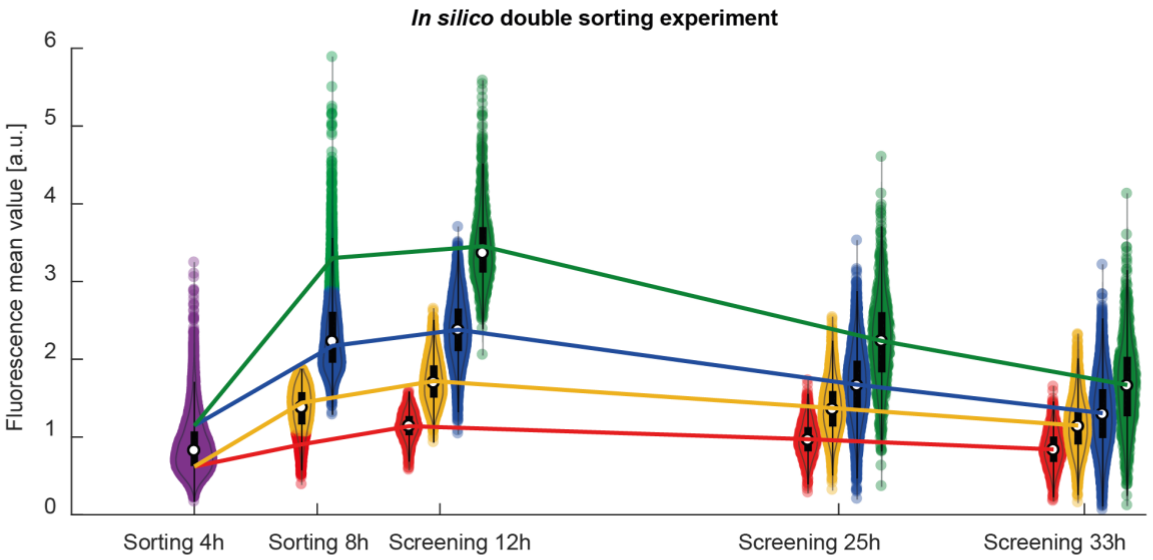


b

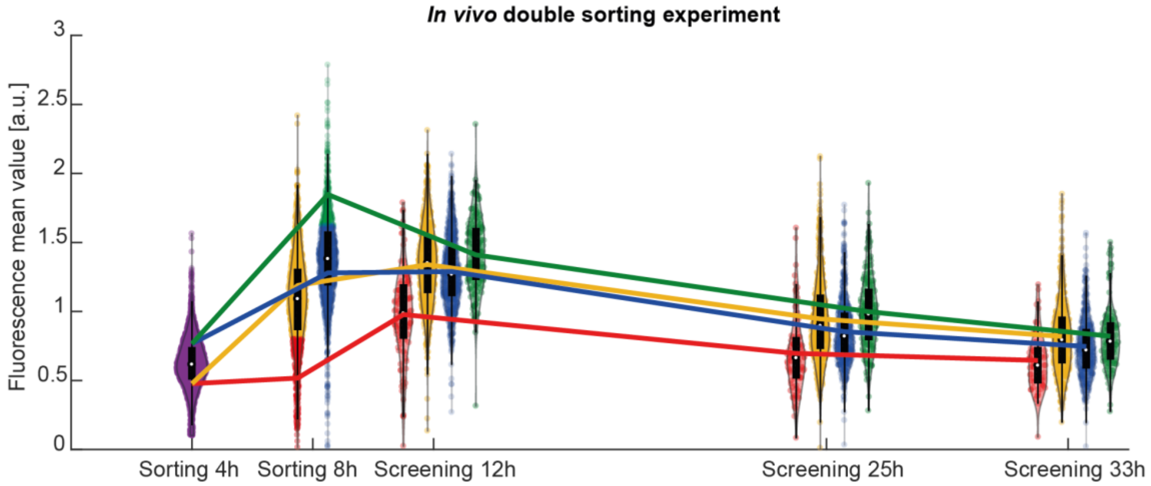


c

| | 8h | | 12 h | | 25 h | | 33 h | |
|---|---|---|---|---|---|---|---|---|
| Group | Mean | CV | Mean | CV | Mean | CV | Mean | CV |
| High-High | 3,30 | 13,6 % | 3,45 | 14,6 % | 2,24 | 26,1% | 1,70 | 34,2 % |
| High-Low | 2,17 | 14,0 % | 2,37 | 16,1 % | 1,67 | 26,8 % | 1,31 | 33,8 % |
| Low- High | 1,44 | 14,1 % | 1,72 | 16,1 % | 1,37 | 23,6% | 1,15 | 28,5 % |
| Low-Low | 0,91 | 14,0 % | 1,14 | 15,2 % | 0,97 | 23,0% | 0,84 | 27,6 % |

d

| | 8h | | 12 h | | 25 h | | 33 h | |
|---|---|---|---|---|---|---|---|---|
| Group | Mean | CV | Mean | CV | Mean | CV | Mean | CV |
| High-High | 1,85 | 11,0% | 1,41 | 18,8 % | 1,00 | 27,4 % | 0,82 | 28,4 % |
| High-Low | 1,28 | 20,3 % | 1,29 | 20,6 % | 0,85 | 27,9 % | 0,75 | 30,5 % |
| Low- High | 1,19 | 22,2 % | 1,34 | 22,0% | 0,95 | 32,8 % | 0,82 | 33,2 % |
| Low-Low | 0,52 | 35,73 % | 0,98 | 34,1 % | 0,70 | 39,9 % | 0,65 | 30,6 % |

**Supplementary Figure 9. Comparison between simulated and measured GFP expression values for double sorting experiment.** Violin plots of the fluorescence mean values of individual *C. elegans* TJ375 simulated (a) and measured (b) in a population double-sorted as described in Figure 2b,c. A comparative chart of the mean and coefficient of variation of this two datasets: *in silico* (c) and *in vivo* (d) is presented as well.

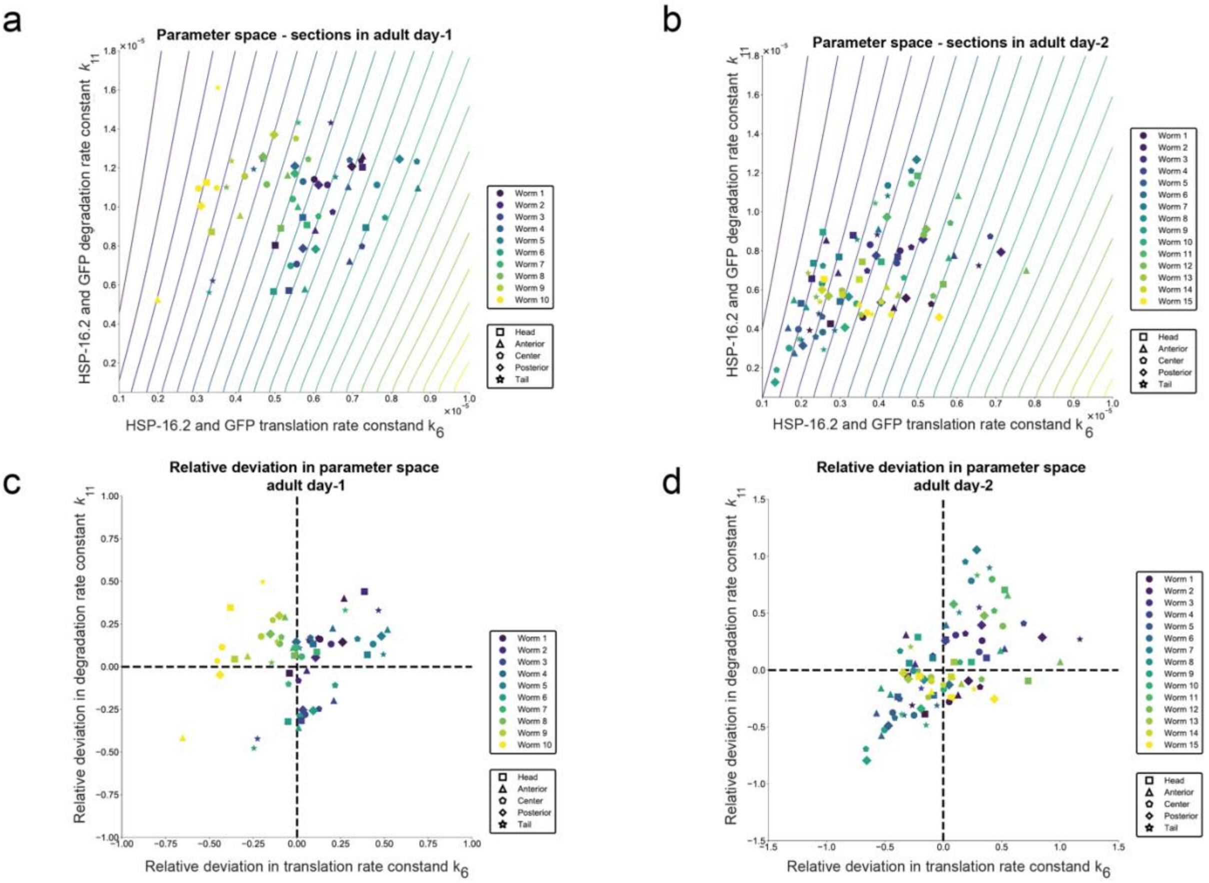


**Supplementary Figure 10. Parameter space analysis of individual tissues of *C. elegans* adult day-1 and day-2.** (a-b) Parameter space ($k_6$ and $k_{11}$) of individual tissues for each *C. elegans* of dataset of adult day-1 (a) and day-2 (b). (c-d) Relative deviation of individual parameters $k_6$ and $k_{11}$ of individual tissues in *C. elegans* of dataset adult day-1 (c) and day-2 (d).

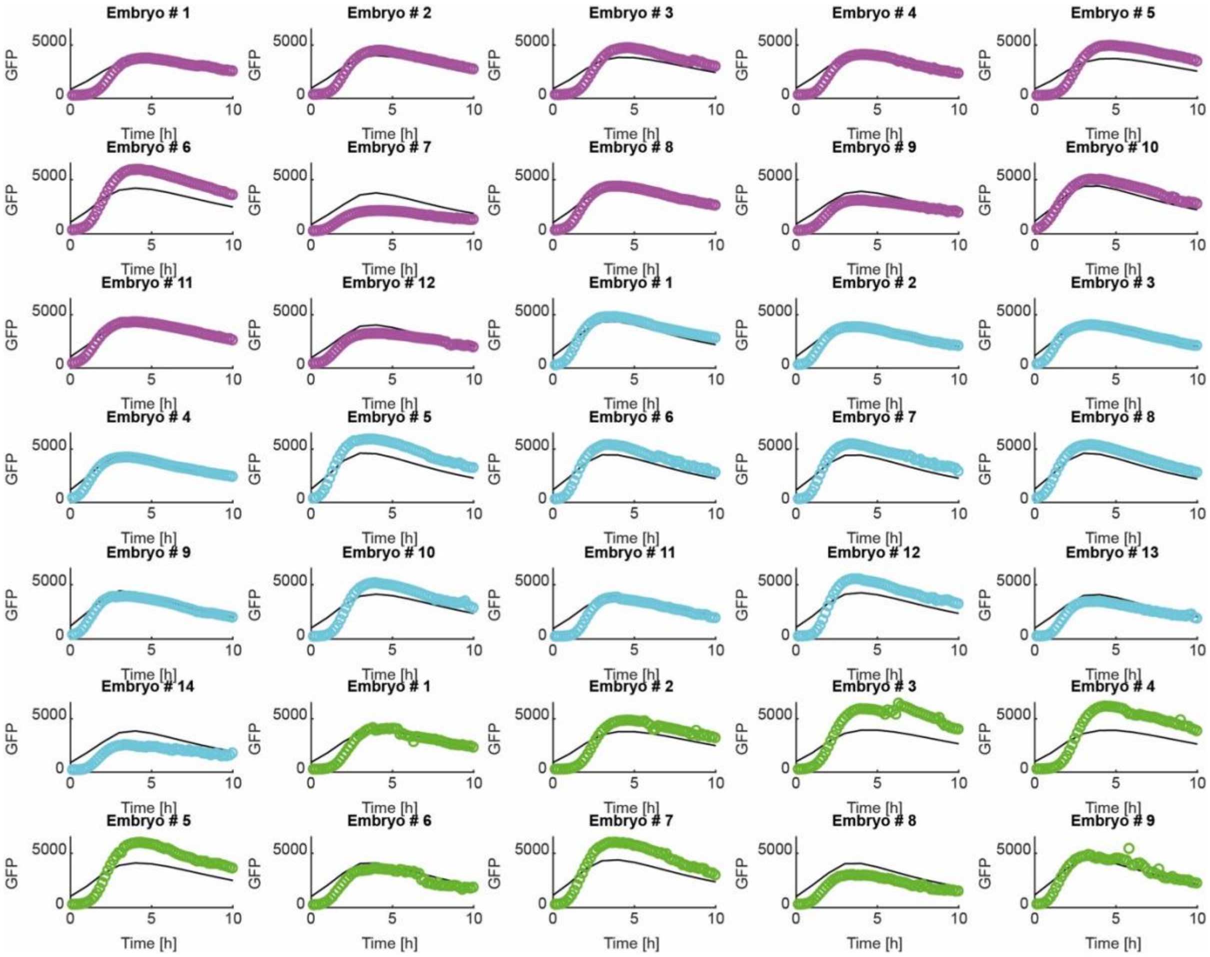


**Supplementary Figure 11. Fitting of individual embryo *C. elegans* HSR with mathematical model.** Individual fitting of individual GFP expression for each analysed embryo shown in Figure 8c, by varying only parameters $k_6$ and $k_{11}$ from the average response. Here the solid lines represent the modelled HSR while the circles represent the experimental data. Circles in magenta represent embryos coming from mothers at day-1 adult, cyan circles embryos coming from mothers at day-2 adult and green circles embryos coming from mothers at day-4 adult.

| | 8 h | | | 12 h | | | 25 h | | | 33 h | | |
|---|---|---|---|---|---|---|---|---|---|---|---|---|
| **Group** | **Mean** | **Std. Dev.** | **CV** | **Mean** | **Std. Dev.** | **CV** | **Mean** | **Std. Dev.** | **CV** | **Mean** | **Std. Dev.** | **CV** |
| **High-High** | 1,85 | 0,20 | 11,0% | 1,41 | 0,27 | 18,8 % | 1,00 | 0,27 | 27,4 % | 0,82 | 0,23 | 28,4 % |
| **High-Low** | 1,28 | 0,26 | 20,3 % | 1,29 | 0,27 | 20,6 % | 0,85 | 0,24 | 27,9 % | 0,75 | 0,23 | 30,5 % |
| **Low-High** | 1,19 | 0,26 | 22,2 % | 1,34 | 0,29 | 22,0% | 0,95 | 0,31 | 32,8 % | 0,82 | 0,27 | 33,2 % |
| **Low-Low** | 0,52 | 0,18 | 35,73 % | 0,98 | 0,33 | 34,1 % | 0,70 | 0,28 | 39,9 % | 0,65 | 0,20 | 30,6 % |

Supplementary Table 1. GFP expression values of sorted sub-populations over time. Comparative chart of the mean and standard deviation of the GFP expression in the resulting 4 groups after sorting at different time points (8, 12, 25 and 33 hours after heat-shock). These fluorescence values were measured with the high-throughput microfluidic system and they are all expressed in a.u.

| **Adult day-1** | | | **Adult day-2** | | |
|---|---|---|---|---|---|
| **Worm #** | **$k_6$ [1/s]** | **$k_{11}$ [1/s]** | **Worm #** | **$k_6$ [1/s]** | **$k_{11}$ [1/s]** |
| 1 | $6.01 \times 10^{-6}$ | $1.14 \times 10^{-5}$ | 1 | $3.58 \times 10^{-6}$ | $4.59 \times 10^{-6}$ |
| 2 | $6.35 \times 10^{-6}$ | $1.11 \times 10^{-5}$ | 2 | $4.54 \times 10^{-6}$ | $8.01 \times 10^{-6}$ |
| 3 | $5.56 \times 10^{-6}$ | $7.07 \times 10^{-6}$ | 3 | $3.77 \times 10^{-6}$ | $8.32 \times 10^{-6}$ |
| 4 | $5.73 \times 10^{-6}$ | $1.13 \times 10^{-5}$ | 4 | $4.46 \times 10^{-6}$ | $7.38 \times 10^{-6}$ |
| 5 | $7.63 \times 10^{-6}$ | $1.11 \times 10^{-5}$ | 5 | $1.93 \times 10^{-6}$ | $3.98 \times 10^{-6}$ |
| 6 | $5.41 \times 10^{-6}$ | $6.99 \times 10^{-6}$ | 6 | $2.55 \times 10^{-6}$ | $3.83 \times 10^{-6}$ |
| 7 | $5.46 \times 10^{-6}$ | $1.04 \times 10^{-5}$ | 7 | $4.22 \times 10^{-6}$ | $1.14 \times 10^{-5}$ |
| 8 | $4.79 \times 10^{-6}$ | $1.12 \times 10^{-5}$ | 8 | $2.53 \times 10^{-6}$ | $6.33 \times 10^{-6}$ |
| 9 | $4.23 \times 10^{-6}$ | $1.16 \times 10^{-5}$ | 9 | $1.69 \times 10^{-6}$ | $3.01 \times 10^{-6}$ |
| 10 | $3.04 \times 10^{-6}$ | $1.09 \times 10^{-5}$ | 10 | $3.42 \times 10^{-6}$ | $5.29 \times 10^{-6}$ |
| **Mean** | **$5.42 \times 10^{-6}$** | **$1.03 \times 10^{-5}$** | 11 | $4.83 \times 10^{-6}$ | $1.14 \times 10^{-5}$ |
| **Std. Dev.** | **$1.23 \times 10^{-6}$** | **$1.76 \times 10^{-6}$** | 12 | $5.15 \times 10^{-6}$ | $8.82 \times 10^{-6}$ |
| **CV** | **22.7 %** | **17.0 %** | 13 | $3.05 \times 10^{-6}$ | $5.96 \times 10^{-6}$ |
| | | | 14 | $3.06 \times 10^{-6}$ | $5.75 \times 10^{-6}$ |
| | | | 15 | $3.69 \times 10^{-6}$ | $4.83 \times 10^{-6}$ |
| | | | **Mean** | **$3.50 \times 10^{-6}$** | **$6.59 \times 10^{-6}$** |
| | | | **Std. Dev.** | **$1.04 \times 10^{-6}$** | **$2.59 \times 10^{-6}$** |
| | | | **CV** | **29.7 %** | **39.3 %** |

Supplementary Table 2. Protein translation and degradation rate constants of individual *C. elegans* day-1 vs day-2 adults. Comparative chart of the calculated protein translation and degradation rates of the *C. elegans* adult day-1 and day-2 studied here.

| Adult day-1 | | Adult day-2 | |
|---|---|---|---|
| **Worm #** | **Slope [a.u./h]** | **Worm #** | **Slope [a.u./h]** |
| 1 | 24.96 | 1 | 12.97 |
| 2 | 26.83 | 2 | 16.79 |
| 3 | 23.34 | 3 | 15.39 |
| 4 | 26.36 | 4 | 12.41 |
| 5 | 39.41 | 5 | 5.77 |
| 6 | 23.32 | 6 | 8.82 |
| 7 | 23.00 | 7 | 11.09 |
| 8 | 18.30 | 8 | 7.74 |
| 9 | 15.35 | 9 | 4.40 |
| 10 | 8.32 | 10 | 11.66 |
| **Mean** | **22.92** | 11 | 11.92 |
| | | 12 | 19.50 |
| | | 13 | 10.31 |
| | | 14 | 8.58 |
| | | 15 | 12.04 |
| | | **Mean** | **11.29** |

Supplementary Table 3. GFP expression slope in *C. elegans* day-1 vs day-2 adults. Comparative chart of the measured GFP expression slopes between the *C. elegans* adult day-1 and day-2 studied here.

| Adult day-1 | | | | Adult day-2 | | | |
|---|---|---|---|---|---|---|---|
| **Tissue** | **Peak [a.u.]** | **Time to Peak [h]** | **Slope [a.u./h]** | **Tissue** | **Peak [a.u.]** | **Time to Peak [h]** | **Slope [a.u./h]** |
| **Head** | 207.11 | 9.7 | 21.4 | **Head** | 120.68 | 10.3 | 11.7 |
| **Int. Anterior** | 219.23 | 10 | 21.9 | **Int. Anterior** | 167.76 | 13.5 | 12.4 |
| **Int. Central** | 234.59 | 8.4 | 27.9 | **Int. Central** | 183.86 | 12.6 | 14.6 |
| **Int. Posterior** | 197.02 | 9 | 21.9 | **Int. Posterior** | 173.71 | 16.7 | 10.4 |
| **Tail** | 136.40 | 10.1 | 13.5 | **Tail** | 140.34 | 16.7 | 8.4 |

Supplementary Table 4. GFP mean expression in distinct tissues in *C. elegans* day-1 vs day-2 adults. Comparative chart of the GFP mean expression in the 5 regions defined in Figure 7a for *C. elegans* adult day-1 and day-2 datasets studied here.

| Adult day-1 | | | | | Adult day-2 | | | | |
|---|---|---|---|---|---|---|---|---|---|
| **Tissue** | **Param.** | **Mean** | **Std. Dev.** | **CV** | **Tissue** | **Param.** | **Mean** | **Std. Dev.** | **CV** |
| **Head** | $k_6$ [1/s] | $5.33 \times 10^{-6}$ | $1.36 \times 10^{-6}$ | 25.5 % | **Head** | $k_6$ [1/s] | $3.33 \times 10^{-6}$ | $1.06 \times 10^{-6}$ | 31.8 % |
| | $k_{11}$ [1/s] | $8.78 \times 10^{-6}$ | $2.03 \times 10^{-6}$ | 23.1 % | | $k_{11}$ [1/s] | $7.21 \times 10^{-6}$ | $1.80 \times 10^{-6}$ | 25.0 % |
| **Int. Anterior** | $k_6$ [1/s] | $5.86 \times 10^{-6}$ | $1.84 \times 10^{-6}$ | 31.4 % | **Int. Anterior** | $k_6$ [1/s] | $3.94 \times 10^{-6}$ | $1.84 \times 10^{-6}$ | 46.7 % |
| | $k_{11}$ [1/s] | $9.29 \times 10^{-6}$ | $2.50 \times 10^{-6}$ | 26.9 % | | $k_{11}$ [1/s] | $6.50 \times 10^{-6}$ | $2.07 \times 10^{-6}$ | 31.8 % |
| **Int. Central** | $k_6$ [1/s] | $6.54 \times 10^{-6}$ | $1.42 \times 10^{-6}$ | 21.7 % | **Int. Central** | $k_6$ [1/s] | $4.14 \times 10^{-6}$ | $1.49 \times 10^{-6}$ | 35.9 % |
| | $k_{11}$ [1/s] | $1.11 \times 10^{-5}$ | $1.80 \times 10^{-6}$ | 16.3 % | | $k_{11}$ [1/s] | $6.35 \times 10^{-6}$ | $2.55 \times 10^{-6}$ | 40.2 % |
| **Int. Posterior** | $k_6$ [1/s] | $5.69 \times 10^{-6}$ | $1.36 \times 10^{-6}$ | 23.8 % | **Int. Posterior** | $k_6$ [1/s] | $3.99 \times 10^{-6}$ | $1.53 \times 10^{-6}$ | 38.3 % |
| | $k_{11}$ [1/s] | $1.12 \times 10^{-5}$ | $1.98 \times 10^{-5}$ | 17.7% | | $k_{11}$ [1/s] | $6.48 \times 10^{-6}$ | $2.87 \times 10^{-6}$ | 44.3 % |
| **Tail** | $k_6$ [1/s] | $4.57 \times 10^{-6}$ | $1.23 \times 10^{-6}$ | 27.0 % | **Tail** | $k_6$ [1/s] | $3.13 \times 10^{-6}$ | $1.23 \times 10^{-6}$ | 39.1 % |
| | $k_{11}$ [1/s] | $1.16 \times 10^{-5}$ | $3.36 \times 10^{-6}$ | 29.0 % | | $k_{11}$ [1/s] | $6.07 \times 10^{-6}$ | $2.60 \times 10^{-6}$ | 42.8 % |

Supplementary Table 5. Adult day-1 and day-2 protein turnover parameters for individual compartments. Comparative chart of the protein turnover parameters for individual compartments in single *C. elegans* for adults day-1 and day-2: mean, standard deviation, and coefficient of variation (CV).

| **Primer name** | **Gene** | **Sequence (5‘ to 3‘)** |
|---|---|---|
| RV70jmjd-3.1fw | *jmjd-3.1* | ggctttctcacgtgcctaat |
| RV71jmjd-3.1rv | *jmjd-3.1* | gctcttttcgtcctgggtata |
| RV72_e1942fw | *ncl-1* | ccgtgatccgaaggttcttg |
| RV73_e1942rv | *ncl-1* | ttccggatccaagcatcatac |
| RV74_e1942seq | *ncl-1* | ccaaatgcccatcttcatcca |

Supplementary Table 6. Sequences of qPCR primers used in this study. qPCR primers used while crossing LSD2088 and LSD2089.

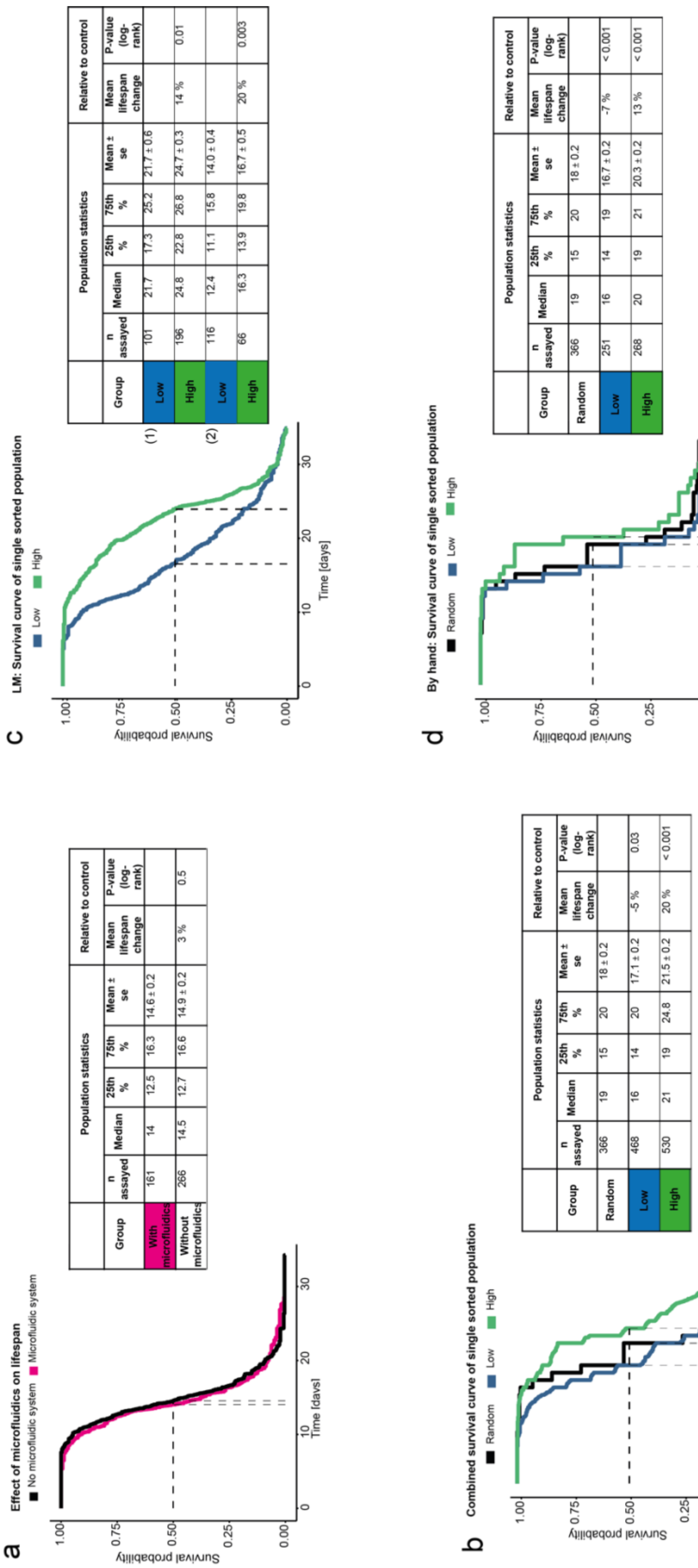


a

| Group | Population statistics | | | | | Relative to control | |
|---|---|---|---|---|---|---|---|
| | n assayed | Median | 25th % | 75th % | Mean ± se | Mean lifespan change | P-value (log-rank) |
| With microfluidics | 161 | 14 | 12.5 | 16.3 | 14.6 ± 0.2 | | |
| Without microfluidics | 266 | 14.5 | 12.7 | 16.6 | 14.9 ± 0.2 | 3 % | 0.5 |

b

| Group | Population statistics | | | | | Relative to control | |
|---|---|---|---|---|---|---|---|
| | n assayed | Median | 25th % | 75th % | Mean ± se | Mean lifespan change | P-value (log-rank) |
| Random | 366 | 19 | 15 | 20 | 18 ± 0.2 | | |
| Low | 468 | 16 | 14 | 20 | 17.1 ± 0.2 | -5 % | 0.03 |
| High | 530 | 21 | 19 | 24.8 | 21.5 ± 0.2 | 20 % | < 0.001 |

c

| | Group | Population statistics | | | | | Relative to control | |
|---|---|---|---|---|---|---|---|---|
| | | n assayed | Median | 25th % | 75th % | Mean ± se | Mean lifespan change | P-value (log-rank) |
| (1) | Low | 101 | 21.7 | 17.3 | 25.2 | 21.7 ± 0.6 | | |
| | High | 196 | 24.8 | 22.8 | 26.8 | 24.7 ± 0.3 | 14 % | 0.01 |
| (2) | Low | 116 | 12.4 | 11.1 | 15.8 | 14.0 ± 0.4 | | |
| | High | 66 | 16.3 | 13.9 | 19.8 | 16.7 ± 0.5 | 20 % | 0.003 |

d

| Group | Population statistics | | | | | Relative to control | |
|---|---|---|---|---|---|---|---|
| | n assayed | Median | 25th % | 75th % | Mean ± se | Mean lifespan change | P-value (log-rank) |
| Random | 366 | 19 | 15 | 20 | 18 ± 0.2 | | |
| Low | 251 | 16 | 14 | 19 | 16.7 ± 0.2 | -7 % | < 0.001 |
| High | 268 | 20 | 19 | 21 | 20.3 ± 0.2 | 13 % | < 0.001 |

**Supplementary Data File 1. Lifespan assays.** Survival curves and statistical analysis of: (a) Effect of microfluidic devices on lifespan. (b) Single sorted animals at 5 hours after heat-shock (3 biological repeats combined, (c) and (d)). (c) Two biological repeats of single sorted animals at 5 hours after heat-shock using the lifespan machine. (d) One by-hand lifespan assay of single sorted animals at 5 hours after heat-shock.

Supplementary Data File 2: Differential equations of our HSR mathematical model

$$\frac{dHSF1}{dt} = -2 \cdot k_{1f} \cdot HSF1_2 + 2 \cdot k_{1r} \cdot HSF1_2 - k_{2f} \cdot HSF1 \cdot HSF1_2 + k_{2r} \cdot HSF1_3 - k_{7f} \cdot HSP \cdot HSF1 + k_{7r} \cdot HSP{:}HSF1 + k_8 \cdot HSF1_2 \cdot HSP + 2 \cdot k_9 \cdot HSF1_3 \cdot HSP + 2 \cdot k_{10} \cdot HSF1_3{:}HSE \cdot HSP$$

$$\frac{dHSF1_2}{dt} = k_{1f} \cdot HSF1_2 - k_{1r} \cdot HSF1_2 - k_{2f} \cdot HSF1 \cdot HSF1_2 + k_{2r} \cdot HSF1_3 - k_8 \cdot HSF1_2 \cdot HSP$$

$$\frac{dHSF1_3}{dt} = k_{2f} \cdot HSF1 \cdot HSF1_2 - k_{2r} \cdot HSF1_3 - k_{3f} \cdot HSF1_3 \cdot HSE + k_{3r} \cdot HSF1_3{:}HSE - k_9 \cdot HSF1_3 \cdot HSP$$

$$\frac{dHSE}{dt} = -k_{3f} \cdot HSF1_3 \cdot HSE + k_{3r} \cdot HSF1_3{:}HSE + k_{10} \cdot HSF1_3{:}HSE \cdot HSP$$

$$\frac{dHSF1_3{:}HSE}{dt} = k_{3f} \cdot HSF1_3 \cdot HSE - k_{3r} \cdot HSF1_3{:}HSE - k_{10} \cdot HSF1_3{:}HSE$$

$$\frac{dmRNA}{dt} = k_4 \cdot HSF1_3{:}HSE - k_5 mRNA$$

$$\frac{dHSP}{dt} = k_6 \cdot mRNA - k_{7f} \cdot HSF1 \cdot HSP + k_{7r} \cdot HSP{:}HSF1 - k_8 \cdot HSF1_2 \cdot HSP - k_9 \cdot HSF1_3 \cdot HSP - k_{10} \cdot HSF1_3{:}HSE \cdot HSP - k_{11} \cdot HSP - k_{13f} \cdot MFP \cdot HSP + k_{13r} \cdot HSP{:}MFP + k_{14} \cdot HSP{:}MFP$$

$$\frac{dHSP{:}HSF1}{dt} = k_{7f} \cdot HSF1 \cdot HSP - k_{7r} \cdot HSP{:}HSF1 + k_8 \cdot HSF1_2 \cdot HSP + k_9 \cdot HSF1_3 \cdot HSP + k_{10} \cdot HSF1_3{:}HSE \cdot HSP$$

$$\frac{dProt_{T=37℃}}{dt} = -\beta \cdot Prot + k_{14} \cdot HSP{:}MFP$$

$$\frac{dProt_{T=20℃}}{dt} = -\alpha \cdot Prot + k_{14} \cdot HSP{:}MFP$$

$$\frac{dMFP_{T=37℃}}{dt} = \beta \cdot Prot - k_{13f} \cdot MFP \cdot HSP + k_{13r} \cdot HSP{:}MFP$$

$$\frac{dMFP_{T=20℃}}{dt} = \alpha \cdot Prot - k_{13f} \cdot MFP \cdot HSP + k_{13r} \cdot HSP{:}MFP$$

$$\frac{dHSP{:}MFP}{dt} = k_{13f} \cdot MFP \cdot HSP - k_{13r} \cdot HSP{:}MFP - k_{14} \cdot HSP{:}MFP$$

$$\frac{dGFP}{dt} = k_6 \cdot mRNA - k_{11} \cdot GFP$$
